# Persistent hippocampal traces of lifelong spatial memory

**DOI:** 10.64898/2026.10.30.762948

**Authors:** Jiang Wu, Markus Meister, Carlos Lois

**Author notes:** these authors contributed equally to this work (Markus Meister, Carlos Lois).

## Abstract

Memory presents a biological paradox: the brain changes continuously, yet a brief experience can influence behavior for a lifetime. How such memories are stored in brain circuits remains unclear. Here we show that two days in a binary-tree maze leave mice with spatial memories that survive repeated 90-day absences over as much as a year. Across return visits, mice explored the maze more efficiently than when naive and resumed direct runs to a reward location sooner, both by a factor of 3. Chronic calcium imaging of the same hippocampal CA1 neurons showed that although most neurons had changed their preferred locations, the population activity at a given maze location still resembled the pattern from 90 days earlier. Furthermore, even when a neuron’s place fields changed location, they tended to remain at the same branching level of the binary tree, reflecting the local geometry of the maze. In return sessions where the neural tuning for branching level was preserved more accurately, the mouse explored the environment more efficiently. Population patterns were best preserved near the reward location, and more reliable neural activity there accompanied earlier direct runs to the reward. With repeated experience, spatial representations changed less, both within a visit and across the long absences. A brief experience thus left selective hippocampal traces that survived months of absence despite drift, grew more durable with each return, and were correlated with memory in navigation behavior.

## Introduction

Animals often have to turn brief encounters with an environment into long-lasting memories. A landscape is organized across scales, from broad terrain and paths to junctions and sites that provide shelter or reward^1,2^. To use such a place again after a long absence, an animal needs to remember where choices lead, which destinations matter, and how to act effectively without complete recall of the layout. However, commonly used navigation tasks can often be solved using local cues, one or two turns, or a beacon at the goal^3,4^. These tasks therefore leave open which features of a complex environment animals remember over long intervals.

The hippocampus is central to spatial navigation in rodents^5^, and it samples inputs from much of the brain^6^, which makes the hippocampus an efficient place to ask what an animal has retained. Many hippocampal neurons are active when an animal occupies particular regions of an environment. These regions are called the neuron’s place fields, and a neuron’s firing pattern across the whole environment forms its place map. These spatial activity patterns have long been considered a candidate neural substrate for spatial memory^7,8^. Yet longitudinal recordings in the same environment show that the place maps of the same hippocampal neurons become less similar over days to weeks^9–13^, and the synapses onto these neurons turn over on similar timescales^14,15^.

Such representational drift and synaptic turnover complicate the understanding of what remains of a spatial memory in the hippocampus^16^. A commonly held view posits that memories can become less dependent on the hippocampus as other circuits support their retrieval^17,18^, but this does not mean that the hippocampal trace has disappeared. Some studies distinguish the survival of a trace from its accessibility: hippocampal ensembles can persist in a silent state, no longer recruited by natural retrieval cues, but recoverable with artificial manipulations^19–22^. Another possibility is that hippocampal stability is selective, such that rather than preserving a detailed place field for every location, the hippocampus may retain only the components of a representation that remain most useful for future navigation^23–26^.

Prior experience can also bias which activity patterns appear when an animal re-enters a familiar place^27–30^. The question is therefore not simply whether hippocampal maps are stable or unstable, but also how the maps evolve when the animal returns, and whether the persistence and rebuilding of brain codes relate to successful navigation.

Most studies of representational drift have recorded the same neurons every day or every few days^9,13,31,32^, so repeated exposure to the task may itself maintain^33^ or update^12^ the representations being measured. Many of these tasks demand little spatial knowledge, so successful performance reveals little about which parts of a representation the animal relies on. To study the persistence of the hippocampal code and its relationship with behavior therefore requires three features: a long interval without exposure to the environment, a complex task that reveals distinct components of spatial memory, and longitudinal recording of the same neurons across the interval.

## Results

### Lifelong memory of a complex navigation task

We let mice solve a maze task in complete darkness across recurring two-day recording cycles separated by approximately 90 days and spanning much of their adult life (Fig. 1a,c,d). The maze was organized as a binary tree (Fig. 1a) embedded in a 15 × 15 grid, with six branch decisions separating the stem from each of 64 end nodes^34^. A fixed water reward was located at one end node and the animal could only get water again from it 90 s after each collection. The maze was connected to the home cage, thus allowing animals to enter, explore, drink, and return home without handling. We refer to the very first day of initial exposure as the naive day and the first day of each later cycle (separated by 90 day gaps) as a return day.

**Figure 1.**
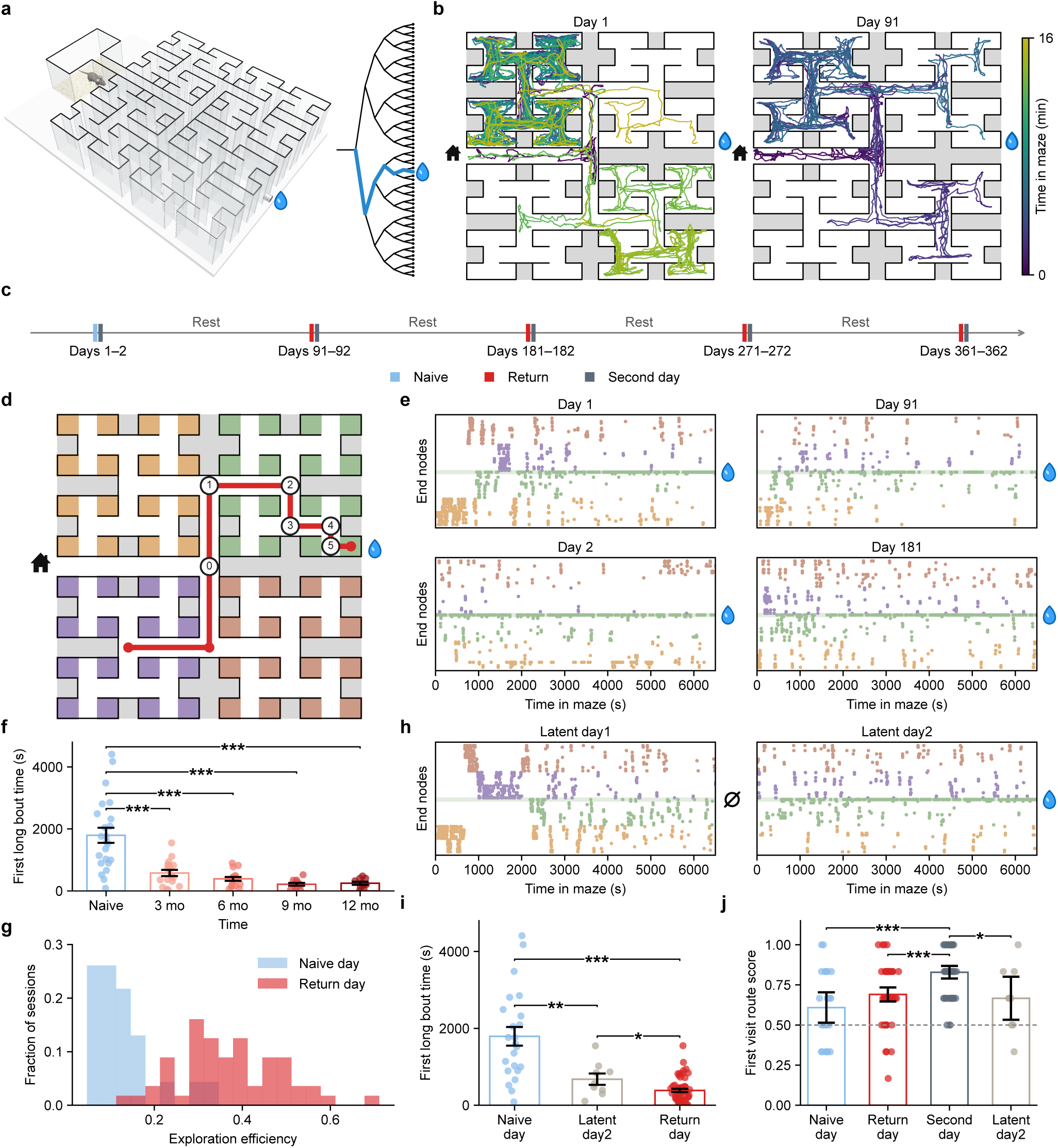
Lifelong memory of a complex navigation task. **a,** The binary-tree maze and its connection to the home cage. The water drop marks the fixed water port. **b,** Trajectories of one mouse from maze entry to its first reward visit on Day 1 and Day 91. Color shows elapsed time in the maze on the same 0–16 min scale. Home and water drop icons mark the entrance and water port. **c,** Longitudinal design of recurring two-day recording cycles. Each cycle comprised two sessions of behavioral and neural recording (Day *i* and Day *i* + 1, *i* = 1, 91, 181, 271, 361), followed by approximately 90 days in the home cage before the next cycle. The Day *i* session was naive in the first cycle and a return session thereafter. **d,** Schematic of the maze, comprising six branch decisions and 64 end nodes in a 15 × 15 grid. The home cage entrance and fixed water port are indicated by the home and water drop icons, respectively. Colored terminal regions define the end node groups used in the end node timelines in **e** and **h**. The red path illustrates an example long bout, defined as a sustained approach to the rewarded end node containing more than six consecutive correct decisions. Circles numbered 0 to 5 mark the six decision nodes scored in **j**. **e,** Example end node visit timelines on Day 1, Day 2, Day 91 and Day 181. The horizontal axis is time spent in the maze (s); the vertical axis identifies the end node visited. Each point marks an end node visit, colored by the terminal region in **d**; the horizontal reference line and water drop indicate the rewarded end node. In this example, compared with naive Day 1, visits on Day 2, 91 or 181 were less clustered from the beginning and concentrated near the rewarded end node earlier in the session. **f,** First long bout time across repeated 3 month cycles. Naive mice required more time in the maze before the first long bout than mice returning after months away. First long bout time fell from 1,794 ± 243 s on the naive day to 580 ± 101, 389 ± 62, 216 ± 43 and 248 ± 50 s after 3, 6, 9 and 12 months, respectively (mean ± s.e.m.; *n* = 23, 17, 17, 12 and 10 sessions, respectively). Each return group differed from naive (regression on log time, with uncertainty adjusted for repeated sessions from the same animal; Benjamini–Hochberg (BH) corrected *p* values were 7.1 × 10^−4^, 1.2 × 10^−6^, 1.2 × 10^−6^ and 2.5 × 10^−5^ at 3, 6, 9 and 12 months). Across return sessions, first long bout time declined with each additional recording cycle (*n* = 56 sessions from 17 animals; fold change per return cycle, 0.72; two-sided *p* = 0.041). Points denote sessions; bars and error bars show mean ± s.e.m. **g,** Distribution of exploration efficiency for naive and return sessions. Return sessions showed less redundant end node sampling than naive sessions. Exploration efficiency was defined as 32/*N*_32_, with *N*_32_ denoting the number of end node visits required to sample the first 32 distinct end nodes during exploration. Efficiency increased from 0.13 ± 0.01 in naive sessions to 0.37 ± 0.02 in return sessions (mean ± s.e.m.; 23 naive sessions from 23 animals and 56 return sessions from 17 animals). Return sessions differed from naive sessions (difference 0.24, 95% CI 0.20 to 0.29; two-sided *p* = 1.1 × 10^−11^). Histograms show the fraction of sessions within each condition. **h,** Example end node visit timelines from a latent control animal. The water port was replaced by a maze wall on latent day1 and restored on latent day2. Point colors follow the terminal end node groups in **d**. At the beginning of latent day1, latently trained mice showed an inefficient exploration pattern similar to that of animals on naive days with reward. They were efficient at the beginning of latent day2. **i,** First long bout time in naive, latent day2 and return sessions. Animals in latent day2 sessions had recent layout familiarity but no prior reward experience, and began the first long bout sooner than naive animals (677 ± 147 s versus 1,794 ± 243 s). Return animals were faster still (385 ± 42 s; mean ± s.e.m.; *n* = 23, 9 and 56 sessions from 23, 9 and 17 animals). BH corrected *p* values were 0.0091 for latent day2 versus naive, 6.1 × 10^−8^ for return versus naive and 0.020 for return versus latent day2. Points denote sessions; bars and error bars show mean ± s.e.m. **j,** First visit route score, defined as the fraction of the six reward path decision nodes at which the animal’s first choice was directed towards reward. Scores are shown for naive, return, second day and latent day2 sessions (*n* = 23, 56, 74 and 9 sessions from 23, 17, 24 and 9 animals). Second day scores exceeded naive, return and latent day2 scores (BH corrected *p* = 4.4 × 10^−4^, 1.6 × 10^−4^ and 0.044). The other comparisons were not significantly different (return versus naive, corrected *p* = 0.16; latent day2 versus naive, corrected *p* = 0.57; return versus latent day2, corrected *p* = 0.74). Points denote sessions; bars show condition means with 95% confidence intervals from regressions with standard errors clustered by animal. The dashed horizontal line marks chance performance (0.5).

On the naive day, mice began by repeatedly revisiting nearby end nodes, then explored broadly for ∼ 1 hour before they started running reliably to the reward (Fig. 1b,e, Extended Data Fig. 1a). To distinguish learning from accidental arrivals at the water port, we measured the time to the first long bout, defined as a trajectory to the water port with more than six consecutive correct decisions. Because the home to water path contains only six decisions, a qualifying long bout had to begin at a remote location within the maze, farther from the water port than the home cage. Over the naive day, mice began completing long bouts regularly and continued to do so on the second day (Extended Data Fig. 1b).

To test whether memory of the task persisted across months, we measured the cumulative time each animal spent in the maze before its first long bout. After the initial 90-day gap, this time fell to roughly one-third of its value on the naive day (Fig. 1f). It decreased further across successive returns at 90-day intervals, up to a year. The number of maze tiles traversed before the first long bout showed the same progression (Extended Data Fig. 1d–f), indicating that the improvement was not explained by faster locomotion. These results showed a durable behavioral improvement, but did not reveal what the animals remembered. The improved performance upon repeated exposures could reflect knowledge of the maze structure, memory of the reward, or a fixed sequence of turns to reward.

If the improved performance was due to knowledge of the maze structure one would expect that mice would waste less time on redundant end node visits. Consistent with this hypothesis, we observed that across return visits, animals sampled the maze with fewer repeated visits to its end nodes than when naive (Fig. 1b,e). We quantified exploration efficiency as 32/*N*_32_, where *N*_32_ is the number of end node visits required to encounter the first 32 unique end nodes. Efficiency was about three times higher on return days than when naive (Fig. 1g, Extended Data Fig. 1h–j). After sampling a branch of the binary tree (Fig. 1a), returning mice were more likely to turn back toward the stem, gaining access to other regions of the maze (Extended Data Fig. 2a–f,i). The increase in stemward turning was greatest near the maze entrance and declined with depth, the number of branch choices separating a junction from the maze entrance (Extended Data Fig. 2g). A biased-walk model incorporating this dependence on depth reproduced most of the exploration gain (Extended Data Fig. 2h).

Mice returning to the maze had experienced both the maze and the reward. To ask what reward experience added to the memory of the maze, we compared the performance of mice that had experienced the maze and the water port with “latent” control animals that had explored the maze without reward on the naive day (Fig. 1h). When reward was introduced, these control mice began long bouts sooner than naive mice. If recent familiarity with the layout was sufficient to explain the return improvement, latent control animals should have performed at least as well as return animals. However, returning mice that had been 90 days away from the maze began long bouts even sooner than latent control mice on their second day (Fig. 1i). This observation is consistent with a learned expectation that the maze offers a reward, which expressed itself as fewer pauses at decision points before the port was rediscovered and as faster approach once it was (Extended Data Fig. 3a–d).

Finally, the improvement in getting rewards could still reflect a memorized fixed sequence of left and right choices, a form of route learning. We tested this possibility by asking whether mice chose the reward-directed branch on their first visit to each of the six decision nodes along the direct path, on second day and after 90-day absence (Fig. 1d,j). The assay detected route learning on the second day of a cycle, but after a 90-day gap returning animals were indistinguishable from naive mice, and their small residual bias was explained by an increased tendency to alternate, which latent controls showed equally (Extended Data Fig. 3e,f). Thus, animals on return days resumed sustained reward runs rapidly, but this behavior did not depend on memory of a fixed motor route.

Brief experience thus left lasting memories of how the maze is structured and of the reward it contains. Both remained evident across repeated absences spanning much of the mice’s adult lives.

### Hippocampal area CA1 keeps a trace after long gaps without maze experience

We next asked whether these memories had corresponding neural traces in hippocampal area CA1 over the same long intervals. In transgenic mice expressing GCaMP6s, we used longitudinal one-photon calcium imaging to track dorsal CA1 neurons across recurring recording cycles. The fields of view remained sufficiently stable to match individual neurons across these gaps (Fig. 2a, Extended Data Fig. 4), yielding 21,412 cross-session neuron matches over 36 session pairs from 14 mice, of which 18,409 met the activity criterion used in the spatial analyses below.

**Figure 2.**
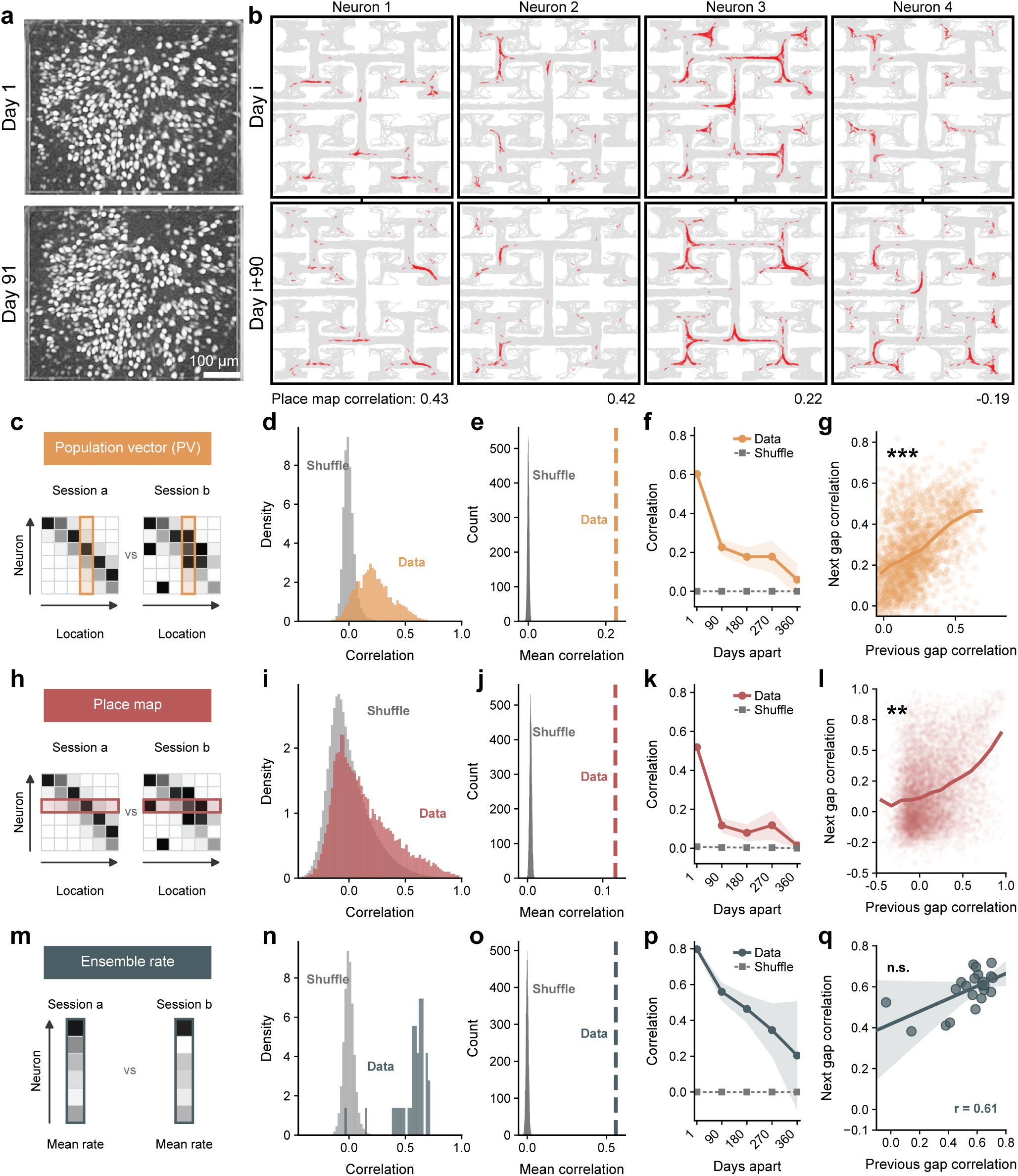
Hippocampal area CA1 retains a persistent trace after long periods without maze experience. **a,** Example of longitudinal CA1 local correlation images from Day 1 and Day 91, showing a stable field of view. Scale bar, 100 *µ*m. **b,** Four example place maps from matched neurons on Day *i* and Day *i* + 90. Each column shows the same matched neuron on Day *i* (top) and Day *i* + 90 (bottom); the value below is the correlation between its paired place maps. The examples span retained and remapped spatial fields, with place map correlations of 0.43, 0.42, 0.22 and −0.19. A red arrow shows the mouse’s position and its orientation when the neuron was active, and the underlying gray lines show animal trajectories. **c, h, m,** Schematics defining population vector (PV), place map and ensemble rate correlations, respectively. Each of these measures compares an array of values between two sessions. PV correlation (**c**) uses the activity at one location across neurons; place map correlation (**h**) uses the activity of one neuron across locations. Ensemble rate correlation (**m**) uses the time-average activity across neurons. **d, i, n,** Distributions of PV, place map and ensemble rate correlations, respectively, for Day *i* to Day *i* + 90 comparisons. The horizontal axes show correlation and the vertical axes show probability density. Each value represents one maze location in a session pair (**d**), one matched neuron in a session pair (**i**), or one session pair (**n**). Colored histograms show the data and gray histograms show neuron ID shuffles. PV correlations showed a broad positive shift; place map correlations were centered near zero but carried a high correlation tail; ensemble rate correlations were strongly positive. The analyses comprised 36 session pairs from 14 animals, yielding 3,810 observations of maze locations, 18,409 matched neuron observations with sufficient calcium events and one ensemble rate correlation per session pair. **e, j, o,** Mean PV, place map and ensemble rate correlations across a 3 month gap compared with 10,000 neuron ID shuffles. Each value in the gray histogram is the mean correlation across session pairs for one shuffle; the colored dashed line marks the observed mean. The horizontal axis shows mean correlation and the vertical axis shows shuffle count. Observed correlations were 0.23, 0.12 and 0.56, respectively, compared with shuffle means of 3.1 × 10^−5^, 0.0041 and 2.1 × 10^−5^ (one-sided permutation *p* = 1.0 × 10^−4^ for each metric). **f, k, p,** PV, place map and ensemble rate correlations across intervals from 1 day to 360 days. The horizontal axis shows the interval between sessions; the vertical axis shows the mean correlation across session pairs, with shaded 95% confidence intervals. Gray dashed lines show neuron ID shuffles. All three correlations declined from adjacent days to month scale gaps but remained above shuffle through 9 months. PV correlations at 1, 90, 180, 270 and 360 days were 0.60, 0.23, 0.18, 0.18 and 0.06. The corresponding place map correlations were 0.52, 0.12, 0.08, 0.12 and 0.01, and ensemble rate correlations were 0.80, 0.56, 0.46, 0.35 and 0.20. The 1 day point summarizes 51 pairs of first and second days within the same cycle from 14 animals. The 90, 180, 270 and 360 day points summarize 36, 22, 11 and 4 pairs of first days from different recording cycles, from 14, 11, 7 and 4 animals, respectively. Comparisons longer than three months included intervening maze visits. **g, l, q,** Recurrence of PV, place map and ensemble rate correlations across successive long gaps. The horizontal axis gives the correlation across one gap and the vertical axis gives the correlation across the next gap for the same maze location (**g**), neuron (**l**), or three-session chain (e.g., days 1, 91, and 181) (**q**). For PVs and place maps, higher correlation in one gap was associated with higher correlation in the next. In **g** and **l**, previous gap correlations were divided into equal width bins; solid curves show the mean next gap correlation at each bin center. Faint points show the underlying 2,266 observations of maze locations and 8,098 matched neuron observations, respectively. In **q**, points show 22 three-session chains (Pearson *r* = 0.61); the line and shading show the fitted regression and its animal-clustered 95% confidence interval. Each analysis included 11 animals with at least three recorded cycles. Regressions of next gap on previous gap correlation, with standard errors clustered by animal, gave slopes of 0.45, 0.35 and 0.31 for PVs, place maps and ensemble rates, respectively. The corresponding BH corrected *p* values were 2.3 × 10^−4^, 0.0020 and 0.063.

To quantify the persistence of memory traces in CA1, we analyzed correlations across time of three types of measurements: population vectors, place maps, and ensemble rates, as illustrated in Fig. 2c, h, m. CA1 population activity at individual maze tiles still resembled the response recorded there 90 days earlier. Across the matched CA1 neurons, the population vector, the pattern of activity across neurons at each location, correlated with its earlier counterpart far more than after shuffling neuron identities (population vector correlation; mean, 0.23; Fig. 2c–e). Yet individual place maps were much less faithful. Some remained strikingly similar across the gap, whereas others changed almost completely (Fig. 2b, Extended Data Fig. 5a). Place map correlations of individual neurons averaged only 0.12, although their mean exceeded the shuffled control and the distribution contained a tail of high correlations absent from the control (Fig. 2h–j).

A neuron could remain one of the more active members of the population even if its place field changes across days. Neurons that were relatively active in one session did tend to remain so 90 days later (ensemble rate correlation; mean, 0.56; Fig. 2m–o). This consistency was stronger in the maze than in the home cage or connecting tunnel (Extended Data Fig. 5e), indicating that the retained signal was related to task engagement rather than merely reflecting stable intrinsic excitability. Because a population vector depends on how active each neuron is at a location, differences in overall activity between neurons that persist over time can contribute to population similarity even when individual place fields change.

Because each recording cycle contained two consecutive days, we next asked whether the long gap persistence depended on which day was compared. All such comparisons spanned approximately 90 days, so any difference between them could not reflect elapsed time. Ensemble rate correlation was highest between the first days of consecutive cycles, indicating that the pattern of relative activity across the population recurred most strongly on re-entry. Population vector correlation followed the same pattern, because this measure also reflects how active each neuron is. However, place map correlation, computed within each neuron and therefore insensitive to its overall rate, was instead similar across all day combinations (Extended Data Fig. 5b–d).

We next asked for how long the CA1 memory trace lasted. We compared the same three stability measures across intervals ranging from 1 day to 3, 6, 9, and 12 months (Fig. 2f,k,p), with comparisons beyond three months including intervening visits. All three measures were highest across adjacent days and declined sharply across month-scale return intervals. Nevertheless, the average correlations remained above the shuffled baseline through the 9-month interval, whereas by 12 months, correlations were no longer distinguishable from chance. Correlations between CA1 activity patterns declined over time, yet remained detectable across intervals of many months.

The wide spread of these three stability measures compared to the shuffled distribution (Fig. 2d,i,n) could arise in two ways. It may reflect a shifting signal, in which different neurons or locations happen to be stable or variable on each return. Alternatively, it may reflect a reproducible organization, in which particular elements of the code are repeatedly more stable than others. The same neurons and maze locations that were most stable across one gap tended to remain so across the next gap (Fig. 2g,l,q). The parts of the representation that changed least therefore tended to be the same from one return to the next.

### Experience makes spatial representations more stable

The preceding analyses compared correlations computed over the entire return days. This was necessary for asking whether any representation survived a months-long gap, but it also compressed the dynamics of the representation that unfolded when an animal re-entered the maze. A whole day correlation could reflect a map that was reinstated immediately at the start of the return day, or it could reflect a representation that was gradually rebuilt during return experience. We therefore measured the time course of map evolution upon returning.

An example neuron makes this ambiguity concrete (Fig. 3a). The neuron recovered a similar spatial pattern after a long absence. However, the place map also changed substantially between the first and second day of each recording cycle. Thus, long gap recurrence and within cycle refinement can coexist in the same neuron.

**Figure 3.**
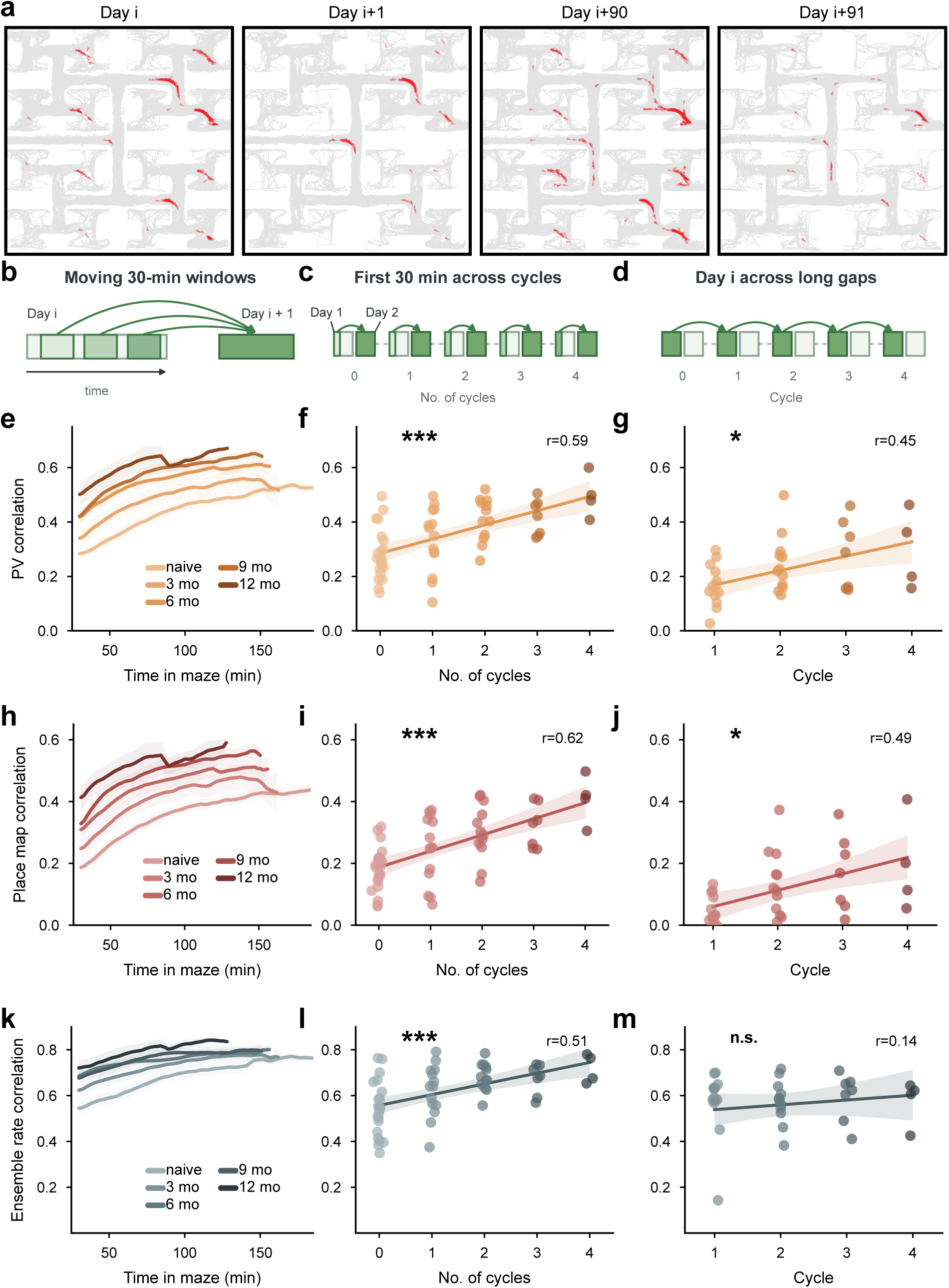
Experience makes spatial representations more stable. **a,** Example place maps from one neuron across two successive recording cycles. Gray lines show animal trajectories and red arrows mark calcium events as in Fig. 2b. Spatial structure recurred between cycles, but the maps also changed between the first and second session within each cycle. **b,** Analysis schematic showing moving 30 min windows from Day *i* compared with the full Day *i* + 1 representation used as the reference. **c,** Representations from the first 30 min of Day *i* were compared with the full Day *i* + 1 reference across recording cycles. **d,** Cross cycle comparison of Day *i* representations from successive recording cycles. The representations were compared between the current cycle and the preceding cycle. **e, h, k,** PV, place map and ensemble rate correlations, respectively, between moving 30 min windows on Day *i* and the full Day *i* + 1 recording from the same cycle. The horizontal axes show time in maze at the end of each window. The point at 30 min summarizes the first 30 min. The vertical axes show the corresponding correlations. Correlations generally increased with time in the maze and began higher after prior experience. Curves show means across animals and shading shows s.e.m. Across **e–m**, darker shades indicate greater cumulative experience. **f, i, l,** Correlation between the first 30 min of Day *i* and the full Day *i* + 1 recording, plotted against recording cycle for PVs, place maps and ensemble rates, respectively. Cycle 0 is the naive cycle; cycles 1–4 are successive returns. Early alignment increased with experience (*n* = 59 observations from 22 animals; Pearson *r* = 0.59, 0.62 and 0.51; BH corrected *p* = 5.5 × 10^−6^, 8.3 × 10^−7^ and 3.4 × 10^−5^, respectively). Each point represents one animal in one recording cycle. Lines are ordinary least squares fits and shading shows 95% confidence intervals. **g, j, m,** PV, place map and ensemble rate correlations between the first days of successive recording cycles, plotted against the current cycle’s number (*n* = 37 session pairs from 14 animals, including one short interval pair). PV and place map correlations increased across returns (Pearson *r* = 0.45 and 0.49; BH corrected *p* = 0.037 and 0.038, respectively), whereas ensemble rate correlation showed no supported trend (*r* = 0.14, corrected *p* = 0.29). Each point represents one pair of sessions. Lines are ordinary least squares fits and shading shows 95% confidence intervals.

To measure this refinement directly, we used the second day as a reference and asked how the naive or return day representation approached that reference over time in moving windows (Fig. 3b). Population vector, place map and ensemble rate correlations all increased as animals spent more time in the maze (Fig. 3e,h,k, Extended Data Fig. 6c,e,g). A comparison of adjacent, nonoverlapping windows showed that the representation changed most rapidly early on the first day and was comparatively stable on the second day, supporting the use of the second day map as the reference (Extended Data Fig. 6d,f,h). Because each cycle produced a different representation, this was not simply recall of one fixed map but a change in how new maps evolved on return.

Prior experience reduced the map difference between the two days of a cycle. Naive visits were most dissimilar from the second day map, whereas return visits began closer to it and remained more closely aligned throughout the day (Fig. 3e,h,k). This improvement was also present when progress was matched by the number of maze tiles visited rather than elapsed time, arguing against a simple sampling explanation (Extended Data Fig. 6c,e,g). An informative way to quantify this improvement across experience is by comparing the representation during the first 30 min of each first day with the full second day (Fig. 3c). The quantification of these early alignments revealed that they increased with cumulative experience for all three measures, confirming our earlier observations (Fig. 3f,i,l).

We next asked whether experience also increased correlations across recording cycles, comparing the first day of each cycle across 90 days (Fig. 3d). Place map and population vector correlations after 90-day gaps increased with cumulative recording cycles, whereas ensemble rate correlation changed little beyond the stability already evident (Fig. 3g,j,m). The maze itself was unchanged, so a fixed response to the same sensory environment could not by itself explain this increase in similarity.

Cumulative experience also increased with age in this longitudinal design. At the same recording cycles, however, neither 1-day nor 3-month correlations increased significantly with age across mice that first entered the maze between 3 and 10 months (Extended Data Fig. 7), arguing against age as the basis of the experience effect.

Together, these analyses showed that both the reorganization within visits and the drift between cycles diminished with repeated experience.

### The reward location favors selective place stability

Across successive returns, both the stability of CA1 representations and the animals’ reward-directed behavior improved in parallel (Fig. 1f and Fig. 3), but these overall trends did not reveal which components of the trace tracked which aspects of performance. The behavioral results revealed two signatures of memory on return: animals resumed long bouts to reward sooner, and they explored the maze more efficiently. We therefore asked whether CA1 carried information specific to either capacity after a long gap.

We trained a spatial decoder on CA1 activity from the first day of one recording cycle and tested it on the first day of the next cycle, 90 days later (Fig. 4a). Its mean error across the maze was 6.0 maze tiles, only modestly below the 6.8 tiles of the control from shuffling neuron identities (Fig. 4b). Thus, the preserved code only weakly supported decoding of maze location, but without much precision.

**Figure 4.**
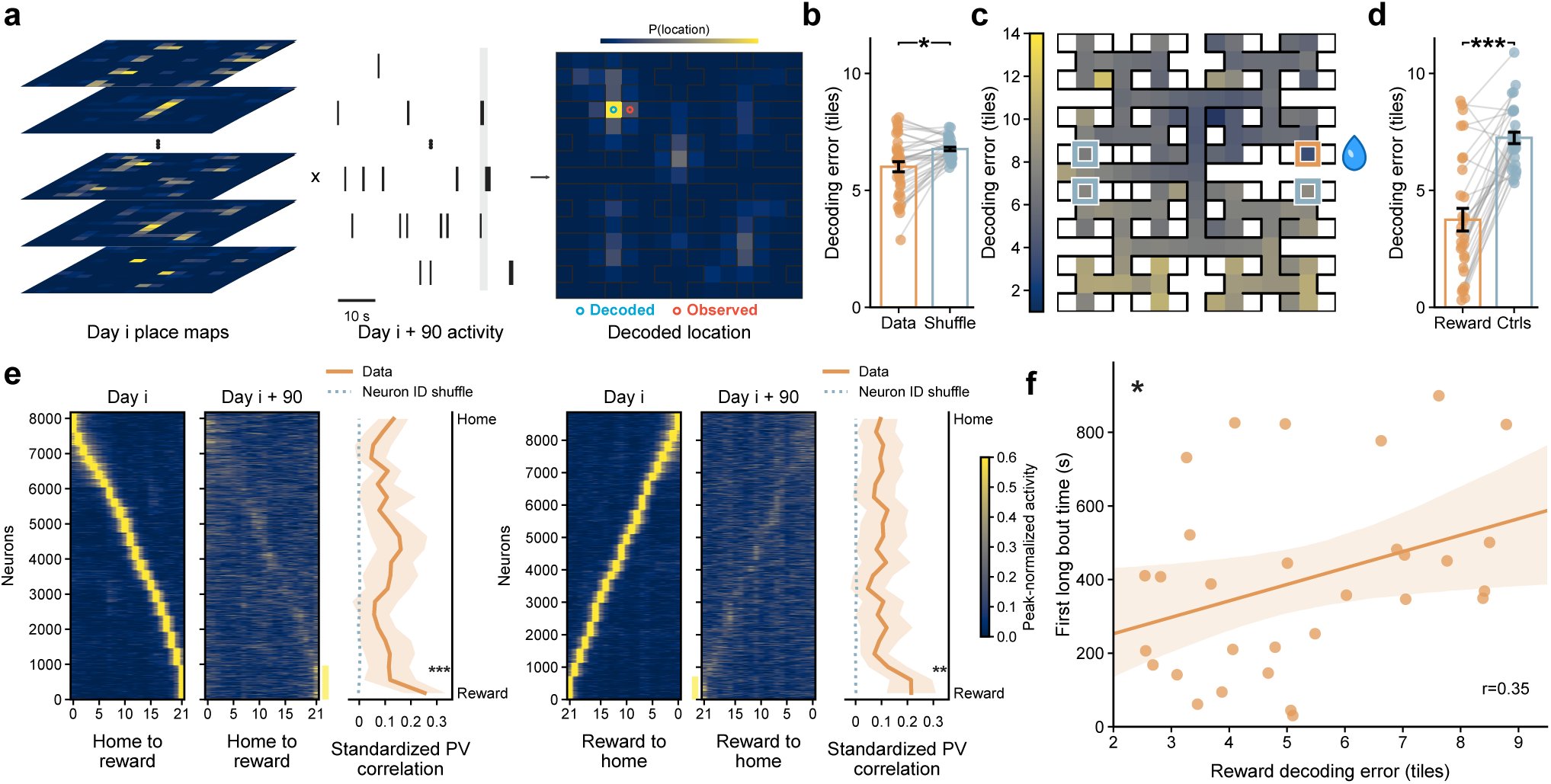
The reward location favors selective place stability. **a,** Schematic of spatial decoding across a 90-day gap. Left, place maps of neurons on Day *i*. Middle, activity of the same neurons on Day *i* + 90, with each tick marking a binarized calcium event. The gray band marks the 1.92 s window ending at the illustrated decoding time. Right, colors illustrate the posterior probability over maze locations; the blue circle marks the decoded location and the red circle the observed location. **b,** Spatial decoding error across sessions, compared with neuron-ID shuffles. Error is the Euclidean distance between decoded and observed locations, measured in maze tiles. Each point is a session pair, and lines connect its observed and shuffled errors; bars and error bars show means ± s.e.m. across pairs. Mean error was 6.0 ± 0.2 tiles for the data and 6.8 ± 0.1 tiles after shuffling (*n* = 36 pairs from 14 animals; two-sided regression with standard errors clustered by animal, *p* = 0.012). **c,** Example spatial map of decoding error for Day 1 to Day 91 session pair. Color gives mean error at each observed maze location; traversable tiles without sufficient samples are left uncolored. The orange square outlines the reward decoder tile, the tile beside the water port used to measure reward decoding error, and blue-gray squares outline three control tiles matched for tree level and local geometry. Errors were 3.4 tiles at the reward decoder tile and 7.0, 7.7 and 8.0 tiles at the controls. **d,** Reward decoding error compared with matched control locations across session pairs whose source Day *i* was rewarded. Error was lower at the reward decoder tile (3.7 ± 0.5 tiles) than at three matched controls (7.2 ± 0.2 tiles). Points denote session pairs, lines connect the reward and control values from the same pair, and bars and error bars show means ± s.e.m. The analysis included 30 session pairs from 14 animals (two-sided regression with standard errors clustered by animal, *p* = 8.6 × 10^−5^). **e,** Population activity along runs from home to the water port (left) and back home (right), compared across 90 days. Each heatmap row is a matched neuron observation from a session pair; columns are route locations. Columns follow the direction of travel, from tile 0 at home to tile 21 at the port on approach, and from tile 21 to tile 0 on departure. Rows are ordered by the Day *i* peak location and retain that order on Day *i* + 90. Heatmaps show peak normalized activity for display. Yellow bars beside the Day *i* + 90 heatmaps mark rows whose peak activity was at the rewarded end node on Day *i*. Standardized PV correlation was plotted horizontally against location vertically, from home at the top to the port at the bottom. For the adjacent correlation profiles, each neuron’s map was standardized separately in each session by subtracting its mean across tiles and dividing by its standard deviation. Positive correlations mean that neurons relatively active or quiet at a given location tended to remain so on both days. Orange curves show means across animals; shading shows 95% confidence intervals; dotted blue-gray curves show the mean of 100 neuron-ID shuffles. At the port, correlation exceeded the mean elsewhere by 0.15 during approach (95% CI, 0.11–0.19; BH-corrected *p* = 4.9 × 10^−4^) and by 0.12 during departure (0.05–0.18; corrected *p* = 0.0026; two-sided exact sign-flip tests across animals). Approach included 8,174 matched neuron observations from 28 session pairs and 13 animals; departure included 8,861 observations from 27 pairs and 14 animals. **f,** First long bout time in the later session plotted against reward decoding error. Each point is one session pair. Lower reward decoding error was associated with an earlier first long bout in the later session (*n* = 30 pairs with rewarded Day *i* source sessions from 14 animals; Pearson *r* = 0.35; two-sided regression of first long bout time on decoding error with standard errors clustered by animal, *p* = 0.022). The line is an ordinary least-squares fit; shading shows a 95% confidence band from bootstrap samples of animals.

The surviving spatial code was not equally reliable throughout the maze. Decoding errors were markedly lower near the reward than at three control locations with matched local geometry (Fig. 4c, Extended Data Fig. 8a). Across all session pairs, errors at the control locations were roughly double those near the reward (Fig. 4d). This difference held after controlling for spatial sampling and the decoder’s spatial prior, and appeared in a population vector correlation analysis as well (Extended Data Fig. 8b,c). The preserved code was therefore strongest near the reward.

Did the same neurons retain their preference for the water port? To separate spatial preference from overall activity, we standardized each matched neuron’s map along the route from home to the port and back. Across the 90-day gap, the population pattern was about twice as similar at the rewarded end node as elsewhere along the route, whether the animal was approaching or leaving it (standardized population vector correlation; Fig. 4e, Extended Data Fig. 8d–i). Shuffling neuron identities completely erased this similarity. At the port, the approach pattern on one side of the gap echoed the departure pattern on the other (Extended Data Fig. 8j). The same neurons thus still marked the reward site months later.

Finally, we asked whether the strength of the preserved code related to behavior. Decoding error across the whole maze predicted how soon mice resumed long bouts, as expected for a behavior that requires traversing a long path (Extended Data Fig. 10a). The relationship held even at the single tile beside the reward, where lower error accompanied an earlier first long bout (Fig. 4f).

Mice explored about three times as efficiently on return as when naive, yet their earlier place maps supported only imprecise location decoding (Fig. 1g and Fig. 4b). What spatial knowledge could support so large an improvement?

### A compact structural code predicts memory guided exploration

Efficient exploration may not require an exact map of every maze location. In the binary-tree maze, a more compact variable is tree level, which groups maze locations at the same depth in the branching hierarchy (Fig. 5a). Tree level is relevant to exploration because the cost of a wrong turn depends on it: re-entering a sampled branch near the leaves costs only a few extra end node visits, whereas doing so near the stem sends the animal back through a large region it has already covered.

**Figure 5.**
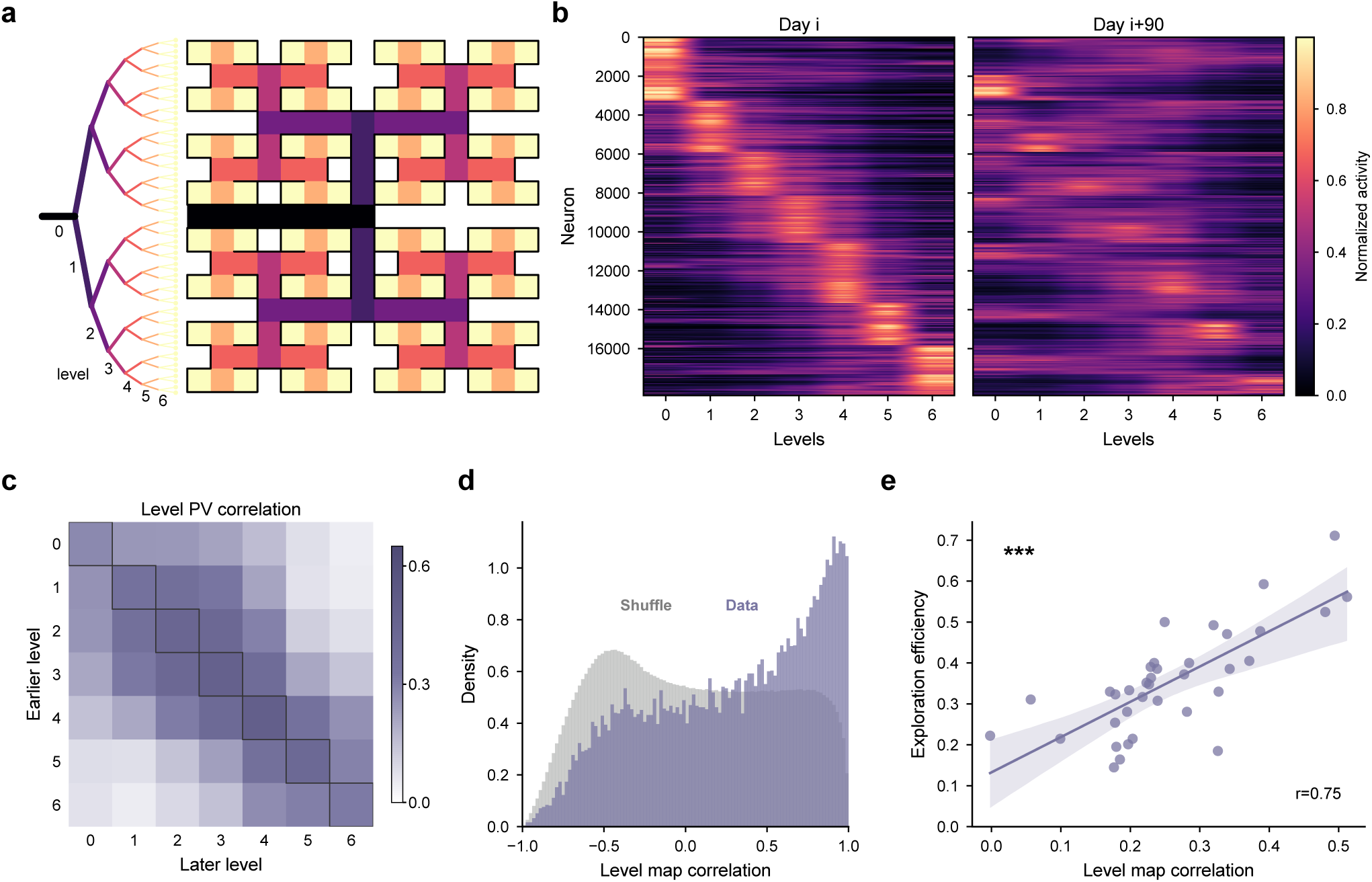
A compact structural code predicts memory guided exploration. **a,** Schematic of the tree level representation. Maze locations were grouped into seven tree levels according to the number of branch choices from the entrance. **b,** Level tuning before and after a 90-day gap. Rows show the same matched neurons in the same order in both sessions; columns show levels 0–6. Color gives each neuron’s normalized activity at that level. Rows are grouped by preferred level on Day *i* (18,409 matched neuron observations from 36 session pairs and 14 animals). **c,** Level PV correlations across the gap. Each cell compares activity across matched neurons at an earlier session level (row) with activity at a later session level (column), averaged across session pairs. Outlined diagonal cells compare the same level; off-diagonal cells compare different levels. Color gives the mean Pearson correlation. **d,** Distribution of level map correlations for individual matched neurons. The purple histogram shows the data and gray shows the neuron ID shuffle; the vertical axis is probability density. The distribution includes a peak near one (mean over neuron observations, 0.28; mean across session pairs, 0.26, as in Extended Data Fig. 9c; *n* = 18,409 matched neuron observations). **e,** Preserved level tuning and exploration efficiency on return. Each point represents one session pair. The horizontal axis gives mean level map correlation across the gap, and the vertical axis gives exploration efficiency in the later session. The line is a linear fit; shading shows a 95% confidence band from 5,000 bootstrap samples of animals (*n* = 36 session pairs from 14 animals; Pearson *r* = 0.75; two-sided regression with standard errors clustered by animal, *p* = 1.7 × 10^−5^).

For each neuron, we computed mean activity at each of the seven tree levels regardless of where in the maze the animals lie (Fig. 5a,b); we call this tuning curve the neuron’s level map. We correlated these level maps across recording cycles to ask whether neurons preserved their tuning to level after a long gap.

As described before, place map correlations were low on average, with most neurons near zero and a long tail of high values (Fig. 2i). In contrast, many neurons retained closely similar level maps, producing a pronounced peak near a correlation of one (Fig. 5d). The mean level map correlation was significantly higher than a shuffle control (Extended Data Fig. 9c), and shifts in preferred level favored nearby levels (Fig. 5c, Extended Data Fig. 9e,f). In the same neurons, level tuning was more stable than tuning to precise location.

Because level maps are coarser than place maps, this stability could reflect averaging activity over fewer groups of locations. We therefore compared them with two similarly coarse spatial maps, each pooling neighboring locations into eight compact regions (Extended Data Fig. 9g). Level maps were more stable than either alternative (Extended Data Fig. 9h).

Level and place coding overlapped without being equivalent. We shuffled locations within each level to break the match between exact locations across days. About half of the mean place-map correlation remained, indicating that the level code carried a substantial part of the spatial similarity across months (Extended Data Fig. 9j). A neuron that preserved its precise field necessarily preserved its level, whereas the converse need not hold. Many neurons retained similar level maps while losing place map correlation, and few showed the reverse (Extended Data Fig. 9i).

This memory of the tree level was also evident in population activity. Activity patterns at the same tree level remained similar across the 90-day gap (Fig. 5c; Extended Data Fig. 9a,b). A decoder trained before the gap recovered the animal’s tree level on return well above chance (Extended Data Fig. 9d). This compact code also recurred across long gaps and became more consistent with repeated experience (Extended Data Fig. 9k–o).

The better the level code was preserved, the more efficiently mice explored on return (*r* = 0.75; Fig. 5e). This single neural measure accounted for more than half of the variation in exploration efficiency (Extended Data Fig. 10b). Compressing the detailed place maps to only a handful of tree levels left their predictive power intact (Extended Data Fig. 10a). Two further comparisons showed that this link was specific. Ensemble rate correlation, although the most persistent of our measures, did not predict exploration efficiency (Extended Data Fig. 10a). Among the turning biases, level map correlation was associated only with the tendency to turn back toward the stem after sampling a branch, the strongest contributor to the modeled exploration gain (Extended Data Fig. 2d, Extended Data Fig. 10c). The value of turning back toward the stem depends on the scale of the maze opened up by that choice. A preserved level code carries information about that scale, and more generally about the affordances associated with the local turning decisions.

Across three months of absence, detailed place maps reorganized, yet CA1 retained the two variables that help solve the maze: where the reward was located, and how much of the maze opened up by a particular choice.

## Discussion

Two days of exploring a maze with 64 dead ends to find water gave mice a memory they could still use 90 days later. On return, mice explored three times as efficiently as when they were naive, and found direct routes to the water port three times sooner (Fig. 1f,g). Detailed tracking revealed a lasting reduction in futile, repeated visits to maze end nodes during exploration, an observation that would be missed by commonly used methods that usually focus on just reaching a goal. This may help explain why rats trained over several days or weeks in earlier studies showed little or no advantage over naive performance a few months later^35,36^. Earlier demonstrations of lasting spatial memory used either simpler spatial tasks, sometimes involving escape from water^37,38^, or months of extensive experience in a complex environment^39^. In our maze, those two days offered repeated opportunities to learn: after finding water, mice could keep exploring at their own pace and come back to it from different places. Such varied experience may have strengthened their memory. Each return after a 90-day absence, gave the mice two more sessions to build on their earlier experience. Across a year of these visits, 90 days apart, they found direct routes to the water port progressively sooner. These gains suggest that each return built on earlier learning. What the mice learned during their first two days could therefore still help them navigate a year later, spanning much of their adult lives.

The hippocampus retained a representation of the maze for months. Many individual CA1 neurons changed where they were active, but overall place maps and population vectors remained more similar than expected by chance even between visits nine months apart (Figs. 2f,k,p). This persistence is striking given the rapid turnover of CA1 dendritic spines^15^. Basal dendritic spines on CA1 neurons have an estimated mean lifetime of about ten days^14^. Neuronal representations drift from one day to the next, but they remain mostly similar. This could account why despite this drift, the behavior does not change across several days. Over months, however, accumulated changes caused by drift could undermine the memories that guide behavior. Tracking the activity of the same neurons for twelve months allowed us to directly investigate what remains of a hippocampal representation after long absences. No previous studies have done this over timescales comparable to this work. Several works have recorded the same neurons every few days for at most two to three weeks, so to be able to compare with our results, we have to extrapolate their reported changes^12,40,41^. Those estimates, together with measurements from works using longer recordings^10,11,42^, suggested that several measures of similarity between visits would decay entirely within one to two months. Therefore, one possibility is that findings from animals repeating a familiar task every day may not generalize to remembering an environment after months away. Another possibility is that our maze requires spatial knowledge that a linear track, the task used in most previous studies, does not. On a linear track, an animal only needs to remember a simple rule: drink from a water port, turn around, and run to the opposite port. Even large changes in how neurons represent locations along the track would not prevent an animal from finding the rewards at its ends. Successful performance of the linear track does not require a memory of the space. Thus, instability in those representations may simply be an indication about how little spatial detail the task demands, rather than what the hippocampus can preserve.^16,43,44^. Across successive returns after 90-day absences, the representations became more stable both within visits and across the months between them (Fig. 3). The maze itself did not change, so its sensory features alone cannot explain why the neurons’ responses became more stable. Renewed experience appears to strengthen the hippocampal memory of the maze. Accounts of preconfigured network dynamics^45,46^ and schemas^47^ proposed such strengthening over shorter timescales, and our results extend it to gaps of three months.

We observed a correlation such that the better neurons preserved their representation of the maze over 90-day absences, the better the mice navigated on return. In fear conditioning studies, even over much shorter intervals of days or weeks, the reported relationships between freezing during recall and the reactivation of neurons involved in learning have been inconsistent^48–51^. In a previous study of mice performing a navigation task every day, most measures of stability of neuronal representations in CA1 were also unrelated to how well they anticipated reward^13^. Our mice explored more efficiently when neurons better preserved their code for how deep the animal was in the branching maze (Fig. 5e). Mice also found direct routes to the water sooner when place maps from an earlier visit more accurately predicted their current location near the reward (Fig. 4f). These strong correlations support the conclusion that mice might actually use their lasting hippocampal representation of the maze to navigate after months away.

The only prior report of memory lasting a year in mice is the so-called “gist memory” after fear conditioning with electric shocks^52^. During remote recall, the animals froze when placed into an unusual context box, regardless of whether that context was associated with shock or safety during training. Hence they only remembered the “gist” of the experience rather than the details. Does the memory documented here similarly refer to the “gist” of the experience, such as the general feel of a plastic maze, rather than its detailed spatial layout? Perhaps the mouse returning after a long gap remembers that such a structure is safe, and thus skips the early period of cautious, tentative cycling, moving right into efficient exploration. Testing this notion would require presenting the animals with a maze with the same dimensions and material, but with a different layout of junctions on return. If the animal was able to navigate this alternative maze efficiently after a long absence, this would represent an interesting case of positive gist memory, in that the mice generalize an attractive response (e.g., the expectation of a reward) to a different context. However, it is unlikely that such gist memory accounts for all the results that we observed in our maze, because we document clear neural memory for specific parts of the maze, and that the quality of that neural trace correlates strongly with the efficiency of exploration (Fig. 5e). No such neural place memory would be needed to convey the general gist of a maze.

What strategies are mice using to remember the maze? A fixed sequence of turns could be memorized to solve this maze. By the second day of every cycle, the mice could be using this strategy, because their first choice at most of the junctions on the direct path already favored the reward branch (Extended Data Fig. 3f). However, after a 90-day gap, that bias was gone (Fig. 1j). This assay samples the first encounter with each junction after return, so it could miss retained route knowledge that required renewed experience to be expressed. Either way, after 90 days, the way mice solved the maze was not directed by a learned sequence of turns. Absences of this length are common for a wild animal, and over such intervals the world does not hold still^53,54^. A learned sequence of turns will fail when a corridor is blocked or if the reward moves. What returned across each gap was not the specific solution to obtain a reward, but the strategies for finding a solution again.

That distinction points to what a durable memory of a place should contain to be useful for an animal. Natural environments confront an animal with the same kinds of situations over and over in different places: a junction where a path divides, a stretch of open ground, a narrowing that leads to shelter. What defines each kind of situation is what it affords rather than where it is. Knowledge of that kind is compact and can be applied in a useful way from one instance to the next^55–59^. Even as the memory of a particular place fades, this compact knowledge would expect to last^60^.

Depth in the branching hierarchy is a variable of this kind: a state abstraction over the many locations that share it. The preserved level tuning could reflect depth itself or one of the features that covary with it in our maze: corridor length, the number of branches ahead, or distance from the entrance. What matters is that such information can guide a choice without a specific memory of the junction: knowing how deep a junction sits is enough to know roughly what a turn will cost, even at a junction not visited for months. Detailed coordinates of location do not generalize this way. Preserving information with precision is costly, and worth keeping only for the places that earn it, such as the reward. At the port, the population code was best preserved whether the mouse was arriving or leaving, and part of that population code was common to both directions, indicating that the mouse held a memory of the location of the reward, and not just about the direction to reach or depart from it (Fig. 4e; Extended Data Fig. 8j). This stable local code close to the reward could help the animal recognize the destination when encountered without specifying the complete sequence of turns needed to reach it.

The paradigms that we used here allowed us to measure which aspects of memory are preserved across a long absence. Remote memory has been studied for decades in relatively simple conditions^18,61^, and tagged c-Fos ensembles have shown that hippocampal neurons can be reactivated upon re-exposure^62–65^. However, knowing that the same neurons participate in learning and recall does not by itself reveal what they represent, whether the layout of a place, an event that happened there, or its familiarity. A representation can preserve some features of a memory while changing others. In a task complex enough to separate components of memory, following the tuning of the same neurons across months makes that selectivity visible. We can foresee three fruitful directions inspired by our observations: what aspects of the information that fades from the hippocampus persist in other circuits, how a changing brain preserves what an animal has learned, and how the brain revises its memory traces when a familiar place presents a new problem.

## Methods

### Experimental procedures

#### Animals

All procedures were approved by Institutional Animal Care and Use Committees at Caltech under protocols 1656 and 1709. Experiments used C57BL/6J-Tg(Thy1-GCaMP6s)GP4.12Dkim/J mice (Jackson Labs stock: 025776)^66^. A total of 36 mice (17 female and 19 male) were used in this study. Of these, 18 contributed at least one return session after a gap of approximately 90 d, and 22 contributed dorsal CA1 calcium imaging, of which 14 had cross-session registration maintained across long gaps. See Supplementary Table 1 for a breakdown of which mice were used in which experiments and analyses. No behavioral or neural measure differed significantly between female and male mice. Mice were 1 to 7 months old at surgery and 3 to 10 months old at their first maze exposure. They were single housed post surgery on a 12 h/ 12 h dark/light cycle and experiments were performed in the dark phase. When mice received water restriction, their body weights were monitored every day. During the intervals between recording cycles (Supplementary Table 3 defines the recurring behavioral and neural terms), mice were housed in a different room and had access to water and food ad libitum.

#### Behavior task

The maze formed a binary tree with six branch choices between the home entrance and each of 64 end nodes (Fig. 1a,d). It was connected to the home cage, allowing mice to enter and leave freely throughout a session. A water port was installed at one end node, which was connected to the entrance by a unique direct route. The rewarded end node was fixed and remained the same across recording cycles. Maze construction followed the previously described design^34^. The apparatus had an approximately 24 × 24-inch footprint and was laid out on a 1.5-inch square grid. Walls were made from 1/8-inch infrared transmitting acrylic (ePlastics) that was opaque in the visible spectrum, and the floor was infrared transmitting acrylic supported by a thicker clear acrylic sheet. The resulting corridors were approximately 1-1/4 inches wide. The walls were 5 inches high and the maze had no ceiling, leaving the top open to accommodate the miniscope tether. Experiments were performed in total darkness under diffuse 940 nm IR illumination (CMVision) for behavior recording. A nose poke at the water port triggered delivery of 20 *µL* water through a steel tube controlled by a Bpod state machine (Sanworks LLC.). After each delivery, the port remained inactive during a 90 s timeout. Recording lasted for 6 h on naive days and 4 h on all the other days. Reward also triggered an infrared LED recorded by the behavioral camera to align with the behavior video. After each recording session, the floor and walls were first wiped with disinfectant wipes (Rescue) and then washed with ethanol and water.

Each recording cycle comprised two sessions that began 24 h apart, Day *i* and Day *i* + 1. Day *i* of the first cycle was the animal’s naive session; subsequent Day *i* sessions were return sessions conducted after approximately 90 d without maze experience (mean ± s.d., 90 ± 4 d; range, 82 to 110 d) with actual interval lengths summarized in Supplementary Table 2. We refer to Day *i* + 1 as the second day of each cycle. Water source (hydrogel) was removed from the home cage 20 h prior to each recording cycle and food pellets were provided on the home cage floor with bedding materials.

In the latent condition, the water port was replaced by a maze wall on latent day1, so mice explored the complete maze without a water port. The wall was removed and a water port was installed later before the start of latent day2, when water became available. The two sessions lasted 6 h and 4 h, respectively. On latent day1, food pellets and hydrogel were provided in the home cage. After the session on latent day1, hydrogel was removed from the home cage. Food pellets remained available between the two latent sessions and during latent day2.

#### Surgery

All surgical procedures were performed in aseptic conditions. Animals were anesthetized in an isoflurane induction chamber at 5% concentration and then placed in a stereotaxic frame on isoflurane at a concentration between 0.5% and 2.5% with body temperature maintained at 37 °C. Ketoprofen (5 mg kg^−1^) and buprenorphine (3.25 mg kg^−1^; Ethiqa XR) were administered subcutaneously before surgery. Bupivacaine (1 mg kg^−1^) was applied along the incision before closure, and ibuprofen was provided in the drinking water for at least 3 d after surgery at 30 mg kg^−1^.

A 1.0 mm or 1.8 mm diameter GRIN lens was implanted on dorsal CA1 for calcium imaging. A 1.0 mm or 1.8 mm diameter craniotomy was first centered 2.0 mm posterior and 1.8 mm lateral to bregma on the right hemisphere. Fresh artificial cerebrospinal fluid was applied throughout the procedure. Using a blunt 26-gauge needle under continuous irrigation, the dura, overlying cortex and part of the corpus callosum were aspirated until a thin layer of horizontal fibers became visible and remained intact over the hippocampus. A GRIN lens (1.8 mm diameter, 0.25 pitch, 0.55 NA, Edmund Optics #64-519 for the custom miniscope and 1.0 mm diameter, 0.5 pitch, 0.5 NA, Inscopix Inc. 1050-002176 for the UCLA miniscope V4) was lowered to 1.35 mm below the skull surface. The lens and two anterior skull screws were secured with cyanoacrylate and acrylic dental cement, and the exposed lens was protected with Kwik-cast.

After at least a month of recovery, mice were anesthetized with isoflurane and a miniscope was aligned to the GRIN lens under live fluorescence to select the focal plane. For animals recorded with the custom miniature microscope, the device was permanently fixed to the implant with dental acrylic and its focusing mechanism was sealed with cyanoacrylate. For animals recorded with the UCLA miniscope, a baseplate was fixed to the implant with dental acrylic and the UCLA miniscope was screwed to the baseplate one day before every recording cycle. The focal plane was relocated each time the UCLA miniscope was reattached using the movie recorded during the previous recording cycle as a template. Mice recovered for at least a week after custom miniscope or baseplate attachment before the first behavior experiments.

#### Calcium and behavioral recording

For 16 animals, hippocampal calcium activity was recorded with a custom miniscope constructed as described previously^10^. The microscope used the sensor board from an MC900 miniature camera containing an OmniVision 7960 CMOS sensor; the sensor board together with the clock and voltage regulator board was connected to a USB analog video to digital converter. LED intensity was controlled through the DAC output of a signal generator. The optical assembly was housed in a black acetal body (Delrin) made by a 5-axis CNC machine (PocketNC) and contained a 470/40 nm excitation filter, a 495 nm long-pass dichroic mirror, a 520/50 nm emission filter (Chroma Technology) and a 12.5 mm achromatic lens (Edmund Optics). The field of view was approximately 600 × 479 *µ*m at 720 × 576 pixels. Another 6 animals were recorded with UCLA miniscope V4^67^.

For the custom miniscope, calcium movies were acquired at 25 Hz using a USB 3.0 video capture device (StarTech.com, USB3HDCAP). For the UCLA miniscope V4, calcium movies were acquired at 25 Hz using Miniscope DAQ. The movies were saved in lossless FFV1 codec format. Excitation power was between 0.05 and 0.2 mW and fixed for each animal and the excitation LEDs were turned off most of the time when animals were outside of the maze. With this precaution, there was no detrimental effect of photobleaching on the optical recordings. A customized miniature commutator system was added between the miniscope and the computer and attached to a pulley system so that the animals could easily turn and travel with the tether.

Behavior was filmed with Grasshopper3 (GS3-U3-23S6M, Flir) and V0828-MPY2 Lens (8 mm, 1.1”, Computar) at 25 Hz and 1,920 × 1,200 pixels. Behavioral and calcium recordings were synchronized at 25 Hz using Bonsai^68^, with a maximum error of 20 ms.

### Data preprocessing

#### Position tracking

DeepLabCut was used to estimate seven body keypoints (nose, tail, body center, and four paws) in each frame using a network trained on approximately 3,000 hand labeled frames^69^. For each recording, four manually annotated maze corners were used to estimate a projective transformation from image coordinates to normalized maze coordinates. Positions were discretized on the 15 × 15 maze tiles. Coordinates with likelihood below 0.95 were removed and linearly interpolated from the remaining detections. Transitions incompatible with the maze topology were repaired between valid anchors using shortest wall respecting paths. Decision node visits were registered upon nose entry into the designated tile of a junction or end node; re-entry into the same node before visiting another node was not counted as a new visit. Nose position was used for route progress and junction choices because it provided the earliest unambiguous entry into a selected branch. Body center was used for whole-maze spatial activity maps and location decoding.

Unless specified otherwise, location dependent neural analyses used speeds greater than 2 and less than 60 cm s^−1^, calculated from nose coordinates. For speed estimates, continuous nose coordinates were median smoothed over five frames within each uninterrupted trajectory. Segments faster than 60 cm s^−1^ were discarded as tracking artifacts. Speed in each behavioral mode was the total nose displacement divided by the time spent in that mode.

#### Calcium imaging preprocessing

Raw calcium videos were transcoded offline to H.264/MPEG-4 AVC in an MP4 container using lossless settings (libx264, preset medium, qp 0, yuv420p). Videos were then temporally downsampled 4× by averaging non-overlapping blocks of 4 consecutive frames, implemented as two successive stages of pairwise frame averaging and decimation. Two recording sessions from the same recording cycle were concatenated and final analysis files were encoded at CRF 10. All procedures were performed in ffmpeg through custom MATLAB (2023b) scripts. Downsampled calcium movies were motion corrected with NoRMCorre^70^ using rigid registration within a manually selected rectangular mask. Shifts were estimated from a spatially band-pass filtered copy of the movie using an expected cell radius of 10 pixels. Motion corrected movies were spatially downsampled 2× for calcium signal extraction.

Local correlation images were used to visualize the CA1 field of view across sessions (Fig. 2a; Extended Data Fig. 4a). Each pixel’s value was the mean temporal Pearson correlation between its fluorescence trace and those of neighboring pixels, following the calculation in CNMF-E^71^. Pixels within an active cell body tend to fluctuate together, making its outline visible.

Spatial components and fluorescence traces were extracted with EXTRACT^72,73^. Extraction used an expected cell radius of 4–6 pixels, spatial high- and low-pass cutoffs of 6 and 1. Components were screened using a trace signal-to-noise ratio greater than 7, a temporal-corruption score below 0.7, a footprint area of 0.1–6 times the expected area, a spatial-corruption score below 5, and an eccentricity below 8. Candidates were then curated with the semi-automated ActSort method^74^: for each movie, human annotation of about 100 candidates was followed by automated classification of the remainder. Fluorescence traces were deconvolved with the OASIS algorithm^75^. Deconvolution used FOOPSI with a first-order autoregressive, AR(1), calcium-dynamics model. Noise was estimated separately for each trace from its high-frequency power spectrum. The decay coefficient was estimated from the trace autocovariance, and the decay and baseline parameters were optimized. The minimum event amplitude was five times the estimated noise standard deviation (*smin* = −5), with no additional *ℓ*_1_ penalty (*λ* = 0). Deconvolution yielded a nonnegative estimate of neuronal activity in arbitrary units. The resulting deconvolved activity was sampled at 6.25 Hz. Each sample was repeated four times to align it with the 25 Hz behavioral record.

#### Longitudinal cell registration

Day *i* and Day *i* + 1 movies processed in a single fit shared component identities by construction. For all other comparisons, spatial footprint projections were first aligned by translation and then refined by nonrigid demons warping^76^. Primary longitudinal identities were constructed from pairwise CellReg registrations^77^ between consecutive recording cycles that were generally separated by approximately 3 months (Extended Data Fig. 4). These intervals typically retained sufficient shared field of view. Pairs for which the two-stage alignment left insufficient overlap for reliable cell matching were excluded from cross-session analyses. Candidate pairs within 14 *µ*m were evaluated using CellReg’s spatial correlation model. Final assignments were made by probabilistic clustering at *P*_same_ *≥* 0.5. Identities spanning 6-, 9-, or 12-month intervals were obtained by transitively composing adjacent cycle maps through their shared intermediate session identities. An identity was retained only when a complete one-to-one chain could be followed through every intervening session.

### Behavioral analysis

#### Behavior mode

Behavioral modes were assigned from the reconstructed node trajectories. We divided maze behavior into reward-directed travel (the “drink” mode), home-directed travel (the “leave” mode), and exploration. Reward-directed travel consisted of direct routes to the water port with all correct turns in rewarded sessions; home-directed travel consisted of direct routes to home with all correct turns. All remaining trajectory segments were classified as exploration.

#### Reward acquisition and locomotor control

We quantified acquisition by when animals first encountered reward and when they began sustained reward-directed navigation (Fig. 1d,f,i; Extended Data Fig. 1a–f). Reward acquisition was described by cumulative reward count. Sustained reward-directed navigation was measured by cumulative long bouts. A node step was movement between adjacent junctions or end nodes in the binary-tree maze. A long bout reached the water port through more than six consecutive node steps, each reducing the shortest path distance to it. First long bout time was the time in maze at the start of the first qualifying bout. A tile step was movement between adjacent tiles of the 15 × 15 maze grid along the reconstructed nose trajectory. We measured progress to the first reward, tenth reward and first long bout in time in maze and tile steps. Cumulative reward and long bout counts were evaluated at intervals of 10 s or 50 tile steps. For statistical comparisons, time in maze and tile steps to each milestone were transformed as log(*x*) because these positive, right-skewed measures differed between groups by several-fold. Regressions against neural measures used time in maze in seconds without a logarithmic transformation.

#### Directed approaches

Reward approaches comprised all long bouts to the rewarded end node that began after the first reward collection. Control approaches also began after the first reward collection and comprised more than six consecutive node steps, each reducing the shortest path distance to the control end node at which the approach ended. The three unrewarded control end nodes occupied locations symmetric to the rewarded end node (Fig. 4c). Eligible control approaches were pooled within session. Reward approach speed advantage (Extended Data Fig. 10a) was defined as

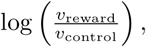

where *v*_reward_ and *v*_control_ were the median reward- and pooled control-approach speeds, respectively.

#### Exploration efficiency

During exploration mode, we measured how much mice revisited end nodes while covering the maze (Fig. 1g; Extended Data Fig. 1h). An arrival at an end node counted as a visit, and a new visit was counted only after the animal left and returned. If *N*_32_ was the visit on which the 32nd unique end node was reached, exploration efficiency was

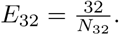

A recorded session had no measured *E*_32_ if it ended before 32 distinct end nodes had been visited. For comparisons of recorded and simulated exploration (Extended Data Fig. 2f,i), such sessions were assigned zero efficiency. Simulated walks that failed to reach 32 distinct end nodes were also assigned zero.

Repeated coverage was measured in coverage rounds ending when 32 distinct end nodes had been visited; counting restarted at the next end node entry, and incomplete final rounds were omitted. Time-resolved round efficiency was summarized in windows of 10 min or 1,000 tile steps (Extended Data Fig. 1i,j). We assigned each completed round to the window in which it ended, averaged round efficiencies within each session and window, and then averaged these session means with equal weight.

#### Junction turning biases and biased walk simulations

We used local turning biases to ask which recurrent choice tendencies were sufficient to reproduce the reduced sampling redundancy of return sessions. Turning biases were calculated from transitions at non-root internal junctions during exploration mode (Extended Data Fig. 2a). After arrival from the stem, *P*_SF_ was the probability of entering either branch instead of reversing. Conditional on continuing into a branch, *P*_SA_ was the probability of choosing right when the current junction lay on its parent’s left branch, or left when it lay on its parent’s right branch. Left and right were viewed from the stem at each junction. After arrival from a branch, *P*_BF_ was the probability of exiting through the stem or sibling branch instead of returning along the incoming branch. Conditional on taking one of these exits, *P*_BS_ was the probability of choosing the stem.

Turning biases were estimated during the first 30 min of time in maze. For analyses comparing all four turning biases and for the corresponding simulations, *P*_SF_ and *P*_SA_ were estimated at each eligible junction and averaged across junctions with equal weight. Because branch arrivals can come from either of a junction’s two branches, *P*_BF_ and *P*_BS_ were estimated separately for each incoming branch, and these branch estimates were averaged with equal weight. For depth-varying *P*_BS_, we pooled decisions within three depth groups: 1–2 (shallow), 3–4 (middle) and 5 (deep). We also calculated a combined depth-weighted stemward probability, giving more weight to choices affecting larger regions of the maze:

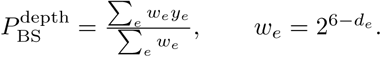

For decision *e*, *y_e_* = 1 if the animal chose the stem and 0 if it chose the sibling branch; *d_e_* is the junction’s depth. The weight is proportional to the number of end nodes below the junction. Analyses comparing all four turning biases included sessions with at least 30 decisions and at least 10 contributing probability estimates for each bias. Depth-varying *P*_BS_ analyses required at least 30 decisions and at least 10 decisions in each depth group.

We simulated walks on the whole binary tree using the measured turning biases. The depth-varying model used separate shallow, middle and deep *P*_BS_ values. At the initial root decision, the two branches were chosen with equal probability. At subsequent junctions, the current and immediately preceding locations identified the incoming edge as a stem or branch. Transitions then followed the corresponding turning bias, and each end node visit was followed by a return to the preceding junction. Each simulated sequence ended after the number of end node visits observed in the corresponding session and was scored with *E*_32_. One hundred walks were averaged for each session.

We asked how much of the observed improvement in exploration efficiency from naive to return sessions could be reproduced by the measured changes in turning biases. For each return session, we simulated exploration using either its own turning biases or those measured when the same animal was naive. We averaged the simulated and observed gains across matched session pairs, then divided the simulated mean by the observed mean. To estimate the contribution of each bias, one return day turning bias was replaced by its value measured when the same animal was naive while the other biases were held fixed. The contribution was calculated as the simulated efficiency before replacement minus the efficiency after replacement.

#### Junction dwell

Junction dwell analyses tested whether prior experience changed the time animals spent at the junctions before and after encountering reward (Extended Data Fig. 3a–d). Dwell was measured for completed choices in which the animal arrived from the stem and exited into a child branch. Each junction was classified as lying on or off the direct home to reward route.

The measurement window began when the animal arrived at the junction and ended when it reached the chosen child node, which could be a junction or an end node. Within this window, dwell was the total time that the nose position occupied the junction tile. Time spent traveling along the connecting corridors was therefore excluded. If the animal left and returned to the junction before reaching the chosen node, that later junction occupancy was also counted. The metric is thus total junction occupancy during a completed choice, not necessarily one uninterrupted dwell. Measurements with dwell greater than 0 s and no more than 30 s were retained.

#### First route choices

At each of the six decision junctions on the direct reward route, we recorded the first exit into either child branch, ignoring any earlier exit towards the stem. First visit route score was the fraction of these six choices directed towards reward (Fig. 1j). Each session therefore contributed one route score, ranging from 0/6 to 6/6. We also examined the first choice at each junction separately, recording whether it was directed towards reward (Extended Data Fig. 3e,f). A session contributed one such value at each of the six junctions.

### Neural analysis

#### Neural activity maps and stability measures

We compared neuronal place maps, population vectors (PVs), ensemble rates and level maps across sessions. We selected neurons using a common threshold set at the 20th percentile of activity rates pooled across neurons and sessions. For this selection, a neuron’s activity rate was its summed unsmoothed deconvolved activity divided by the duration. Primary place map, PV and level map comparisons, and decoding analyses, required the threshold to be met in both sessions. Ensemble rate analyses used the inclusion rules stated below. Unless stated otherwise, we smoothed deconvolved activity in time with a Gaussian kernel of 1 s half-width at half-maximum and used it in all the following analyses. Smoothing was applied within uninterrupted recording segments before selecting analysis frames based on animal speed. Each place map summarized a neuron’s activity at each location:

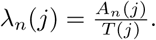

Here, *A_n_*(*j*) is the summed activity of neuron *n* at maze location *j*, and *T* (*j*) is the occupancy time in seconds. The map therefore describes average activity. All quantitative spatial analyses used the 15 × 15 maze grid. Selected place maps were rendered on a 45 × 45 grid only to visualize finer spatial structure. When comparing the same location (or level) across sessions, we required at least 5 s of occupancy in each session. Primary maps were not spatially smoothed, so activity was not mixed across maze walls. Spatial smoothing was applied only to the transformed maps in the within-level shuffle control described below.

In the resulting activity matrix, rows represent neurons and columns represent locations (Fig. 2c,h). PV correlations compare columns across sessions; place map correlations compare rows. Superscripts *A* and *B* below identify the two sessions being compared. We used Pearson correlation throughout; the subscript on corr identifies the dimension over which it was calculated.

To ask whether a location evoked a similar population response in the two sessions, we correlated activity across the matched neurons:

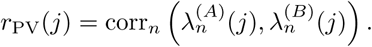

We averaged these correlations across the maze locations sampled in both sessions to obtain one PV correlation.

To ask whether an individual neuron retained its spatial activity pattern, we correlated its maps across locations:

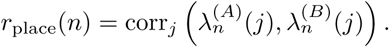

We then averaged the neuron specific correlations within each session pair.

For the primary level map, we replaced each sample’s spatial location with its tree level (Fig. 5a,b):

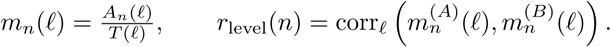

Here, *A_n_*(*ℓ*) and *T* (*ℓ*) are the summed activity and occupancy time pooled across locations at level *ℓ*. The primary level maps used levels 0–6. Each sample’s level was that of the maze tile containing the body center. A sample whose body center fell outside the mapped maze tiles was assigned the level of the most recent maze tile, and samples before the first maze tile of a session had no level and were excluded from level maps.

Ensemble rate correlation asked whether neurons that were relatively active in one session remained so in the other. For each neuron, we summed its activity and divided by the total duration, giving the session-wide value *λ̅_n_*. We correlated these means across matched neurons:

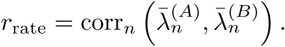

The bars denote time averaged activity, with duration calculated from the retained samples. Ensemble rate correlations retained matched neurons that were silent in one session.

#### Long gap correlation and recurrence

We grouped session pairs by elapsed time. The one-day group compared the first and second days within a recording cycle, whereas longer intervals compared first days from different cycles (Fig. 2f,k,p; Extended Data Fig. 9k).

To test whether stability across one long gap predicted stability across the next, we regressed later gap correlations on preceding gap correlations in three-session chains (Fig. 2g,l,q; Extended Data Fig. 9l). Place and level map analyses followed neurons matched across all three sessions and required the common activity threshold in all three. Each neuron contributed one pair of correlations.

PV recurrence used the same three-session matching and activity threshold requirement. Each location with sufficient samples across both gaps contributed one pair of correlations. Ensemble rate recurrence used each gap’s own matched population, retaining neurons silent in one session, and each chain contributed one pair of correlations. Regressions used these individual observations; binned means were used only for display.

#### Ensemble rate correlation during and outside maze behavior

We compared ensemble rate correlation during maze behavior with correlation outside the maze using Day *i* to Day *i* + 90 pairs (Extended Data Fig. 5e). Outside maze samples combined all frames recorded in the home cage or connecting tunnel. Within each session pair, we drew the same number of frames from inside the maze as from outside it. The inside-maze sample included stationary periods as well as movement, matching the absence of speed selection outside the maze. Ensemble rate correlations were averaged across 20 draws.

#### Within cycle changes in neural representations

We measured how similarity to the full Day *i* + 1 representation varied over the course of Day *i* (Fig. 3b,e,h,k; Extended Data Fig. 6a,c,e,g; Extended Data Fig. 9m). Place maps, PVs and ensemble rates were calculated in overlapping 10, 20 or 30 min Day *i* windows advanced by 60 s and in windows of 3,000 tile steps advanced by 100 tile steps. Level maps were calculated in overlapping 30 min windows advanced by 60 s. Each representation was correlated with the corresponding full Day *i* + 1 representation. Each correlation was plotted at the end of its window, in time in maze or maze tile steps. Time windows were included when they contained at least 300 s of retained calcium samples.

For each window definition, *N*_w_ was the number of nonoverlapping windows of that span that fit within the session, calculated from time in maze or maze tile steps as appropriate. A spatial location required at least 5/*N*_w_ s in the Day *i* window and 5 s in the Day *i* + 1 session to be considered in the analysis. Correlations were first calculated within recording cycle and then averaged with equal weight across animals for group curves.

To measure local stability within a day, we correlated the representations in the 30 min before and after a sliding pivot (Extended Data Fig. 6b,d,f,h). Pivots were advanced by 60 s. Place map and PV comparisons used neurons that met the common activity threshold over that day, whereas ensemble rate comparisons used all neurons. Within each recording cycle, Day *i* and Day *i* + 1 were restricted to pivot positions valid in both sessions, beginning at 30 min and ending at that cycle’s last shared pivot. Each window side required at least 300 s of included calcium data.

#### Cross-session location decoding

To test whether a spatial code persisted across sessions, we trained a decoder on the earlier session of each pair and used it to estimate the animal’s location in the later session (Fig. 4a–c; Extended Data Fig. 8a).

We first converted the unsmoothed deconvolved activity into a binary sequence. Then, for each decoded frame *t*, the input *q_tn_* was the number of ones in that sequence for neuron *n* in the 48-frame window ending at *t*. From the training session, we estimated the expected count *µ_n_*(*j*) as the mean of these counts at location *j*.

Let **q***_t_* be the vector of counts across matched neurons. We approximated these counts as independent Poisson variables given location^9,78^,

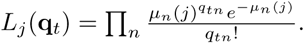

Let *π_j_* be the fraction of retained training time at location *j*. We combined this occupancy prior with the likelihood and selected the location with the largest posterior probability,

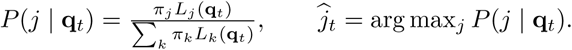

The decoding readouts related to behavior in Fig. 4f and Extended Data Fig. 10 instead used a uniform prior, with equal *π_j_* for all locations.

Location decoding error was the Euclidean distance on the maze grid between the decoded and observed locations of the animal’s body center. We expressed it in maze tiles and averaged it over the eligible target frames of each session pair.

For the neuron-ID shuffle, registered neuron IDs in Day *i* + 90 were permuted relative to the fixed Day *i* expected-count templates. The same neurons and target frames were retained, and the decoder was not refitted. Error was averaged across 1,000 shuffles within each session pair.

For an individual error map, errors were averaged within observed location, and locations with fewer than five test frames were left blank. For the cohort spatial error map, location specific errors were averaged within session pair, within animal and across animals, giving each animal equal weight. Only locations represented in every contributing animal entered the mean error map.

#### Reward code stability

Reward decoding error was evaluated at the tile immediately next to the rewarded end node and at three nonreward tiles matched for tree level and local geometry (Fig. 4d). Analyses using Day *i* as the source were restricted to pairs in which both sessions were rewarded; pairs with an unrewarded source session were excluded. All otherwise eligible Day *i* + 1 source sessions met this criterion. The primary analysis used all retained test frames at these locations and the empirical training occupancy prior.

To test whether unequal spatial sampling or the empirical occupancy prior explained the lower decoding error near reward, we balanced frames or visits across the reward and qualifying control locations (Extended Data Fig. 8b). Locations entered these comparisons when they had at least 20 retained frames in both source and target sessions. Pairs required the reward location and at least one qualifying comparison control location. For frame balancing, each draw sampled the same number of frames at every qualifying location, with the number set by the least-sampled location in each session. The source and target sessions were balanced independently. A visit comprised consecutive retained samples from the same location separated by no more than five acquisition frames. For visit balancing, we averaged source decoder counts and target decoding error within visits, so each sampled visit contributed equally regardless of its duration. We then sampled equal numbers of visits across locations, again setting the count independently in each session. In each draw, we re-estimated source expected-count templates only at the compared reward and control locations, using sampled frames or equally weighted visit means. Expected counts at all other locations were unchanged. Both controls retained the full-maze decoder, with uniform prior weights across training-supported locations, and used 1,000 resamples. Resamples were averaged within session pair and then within animal.

We next asked whether PVs at the reward location were more similar across the two sessions in each pair than PVs at the control locations (Extended Data Fig. 8c). For reward targets, same location comparisons correlated the reward PV with its earlier reward location vector; cross location comparisons averaged its correlations with the three earlier control location vectors. For control targets, same location comparisons correlated each of the three control vectors with its earlier counterpart and averaged the three correlations; cross location comparisons correlated each control vector with the earlier reward location vector and averaged the three correlations.

#### Place maps along linearized routes

Home to water runs followed the unique direct route between the home and the water port; water to home runs followed the reverse route. We also analyzed routes connecting the water port to the junctions labeled node 2 and node 3 in Extended Data Fig. 8h, in both directions. All route comparisons used pairs whose Day *i* session was rewarded. The primary comparison used full-session maps from Day *i* and Day *i* + 90, separately for approach to and departure from the port. Each map described mean unsmoothed activity at each nose position along the route. Both sessions required at least five runs in the analyzed direction. A tile was considered observed when it contained at least 0.2 s of retained samples.

To compare relative spatial preferences, we standardized each neuron’s map separately in each session. If *a_n_*(*x*) is the mean activity of neuron *n* at route location *x*, then

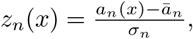

where *a̅_n_* and *σ_n_* are the mean and standard deviation across tiles. A positive value of *z* indicates activity above that neuron’s average across the route. Tiles were weighted equally; when support was incomplete, only tiles observed in both maps were used. At each location, we computed the Pearson correlation across matched neurons,

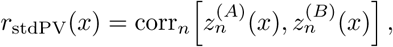

where *A* and *B* label the two maps being compared. We call this the standardized PV correlation (Fig. 4e; Extended Data Fig. 8d–j). The port advantage was the correlation at the rewarded end node minus the mean correlation across the remaining tiles. Neuron-ID shuffles permuted the later session’s neuron identities while preserving its maps (100 permutations per session pair).

At the rewarded end node, approach maps included moving frames within 0.5 s of entry, and departure maps included moving frames within 0.5 s before exit. A gap longer than 0.4 s between moving frames ended the window, assessed forward from entry for approach and backward from the first moving frame at or after exit for departure. Frames claimed by both directions were assigned according to nose movement along the route; frames without clear direction were excluded.

To test whether the port advantage reflected more reproducible activity within a day, we divided each session’s runs into odd and even runs in temporal order. All four maps were restricted to the same observed tiles before standardization. Across-gap correlation was the mean of the four pairings between half-session maps. Within-day correlation compared odd and even runs separately in each session. We averaged each of these three quantities within each animal, then divided the across-gap mean by the geometric mean of the two within-day means. For the port, averages were taken across session pairs; for elsewhere, they also included the other tiles.

For the equal frame control, we sampled the same number of frames in each of the tiles on the linearized routes. This number was each session’s smallest retained frame count across all tiles. Maps and standardized PV correlations were recomputed in 100 draws before averaging within session pair. Neurons selected from the full-session maps were retained when inactive in a half-session or resampled map, with their standardized values set to zero for that map.

For the reward delivery control, home to water runs were classified as rewarded if water was delivered within 3 s of entry into the rewarded end node, and as timeout otherwise. Maps were calculated separately for the two run types, using neurons active in both types and sessions.

The second day of each visit was analyzed in the same way, comparing Day *i* + 1 with Day *i* + 91 and selecting neurons on those recordings. Neuron matching was inherited from the jointly extracted recordings within each recording cycle. For cross-direction comparisons, we used session pairs and matched neurons available in both directions. Maps were aligned by physical location; we correlated approach on Day *i* with departure on Day *i* + 90, and departure on Day *i* with approach on Day *i* + 90, then averaged the two correlations.

#### Level maps and controls

For level PV correlations, each level was represented by a vector containing the activity of matched neurons that passed the activity rate threshold. We computed the Pearson correlation between every earlier-session and later-session level vector to form a seven-by-seven matrix (Fig. 5c). Its diagonal compares the same level across sessions; the remaining entries compare different levels. Only levels with at least 5 s of occupancy in both sessions were included. Each displayed matrix entry is a mean across eligible session pairs, whereas the overall same-level correlation was averaged first across levels within a pair and then across pairs. Neuron ID controls used 10,000 later-session identity permutations, shared across levels within each repeat. These correlations used the original activity maps, before display normalization.

To decode tree level, we applied the location decoder’s 48-frame count definition and Poisson model to the seven primary levels rather than to maze locations (Extended Data Fig. 9d). Each level required at least 40 s of training samples. Accuracy was the fraction of test frames assigned to the correct level. The neuron-ID control permuted later-session identities relative to the fixed earlier-session expected counts without refitting, and accuracy was averaged over 200 shuffles per session pair.

A neuron’s preferred level was the unique maximum of its level map. Maps that were absent, constant, tied at the maximum or insufficiently occupied had no preference. Among neurons whose preferred level differed between sessions, we calculated the probability of each later preferred level given the earlier one (Extended Data Fig. 9e,f). Probabilities were normalized within each earlier level. The null distribution was obtained by permuting the later session’s preferred levels within session pair 10,000 times and identifying neurons with changed preferences anew after each permutation.

To disrupt correspondence between exact locations across sessions, we rearranged eligible connected groups of maze tiles in the later-session place maps, keeping each group within its tree level (Extended Data Fig. 9j). We repeated this rearrangement 1,000 times, using the same permutation for every neuron in each repeat. Each neuron’s shuffled map was then smoothed once across adjacent tiles at group boundaries to reduce discontinuities introduced by the rearrangement. We then recalculated place map correlations between the shuffled later-session maps and the original earlier-session maps.

To test whether averaging locations alone increased stability, we compared the level maps with two coarse spatial partitions (Extended Data Fig. 9g,h). We removed the connecting corridors, divided the remaining maze into sixteen square blocks, and merged adjacent block pairs horizontally in one partition and vertically in the other, giving eight regions in each. Each region required at least 5 s of data in each session.

#### Neural–behavior associations

##### Primary associations

First long bout time in the later session was regressed on reward decoding error from a decoder trained on the rewarded Day *i* source session (Fig. 4f). Both variables were standardized. Neurons were registered across Day *i*, the paired Day *i* + 1 source and the later target session. Ensemble size was set by the smallest qualifying shared population across the included session pairs, allowing every comparison to use an equally sized population. Neurons were sampled in 50 random draws, while one fixed 80 s sample of reward location activity from the target session was held constant. These decoders used a uniform prior over candidate locations.

For the primary structural code association, we regressed exploration efficiency, *E*_32_, in the later session on mean level map correlation across the preceding gap. Each successive pair of Day *i* sessions separated by approximately three months contributed one observation (Fig. 5e).

##### Cross measure comparison from matched population

To compare the behavioral associations of the six neural readouts on equal terms, we used successive long gap session pairs in which all six readouts and the relevant behavior from later sessions were defined (Extended Data Fig. 10a). Within this common set, reward decoding error, whole-maze decoding error, place map correlation, level map correlation, PV correlation and ensemble rate correlation were recalculated from coordinated, equally sized samples of registered neurons. Population size was set by the smallest qualifying registered population across these pairs. Neurons were sampled in 50 random draws, and the same identities were used for all six measurements within a draw. Each readout was averaged across draws within its session pair before it was related to behavior. Both decoders used a uniform prior over candidate locations. Reward decoding used 80 s of test activity from the reward location in every pair. Whole-maze decoding used 80 s sampled across the maze without balancing occupancy. The number of spatial locations was set by the session pair with the fewest source–target locations in common; each pair sampled this number from its own common locations. For the matched population level maps, we averaged the sampled location means within each level from 0 to 5. Level 6 was excluded because it was not represented in four included session pairs. For ensemble rate, 80,000 frames were equally sampled from each session.

##### Commonality analysis

We used commonality analysis to ask whether reward decoding and level map correlation explained overlapping or distinct variation in behavior (Extended Data Fig. 10b). We fitted first long bout time and exploration efficiency separately, using each readout alone and both together. Let *R*^2^_reward_, *R*^2^_level_, *R*^2^_both_ denote the fractions of outcome variance explained by these three models. Each readout’s unique contribution was the gain in explained variance when it was added to the model containing the other. The shared component was the remainder after subtracting both unique contributions from the joint model’s explained variance:

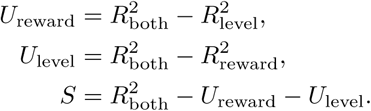

The three components sum to the variance explained by the joint model. All models were ordinary least-squares fits with an intercept, using the same session pairs and behavioral outcome; *R*^2^ was evaluated on those pairs.

We also tested whether reward decoding and level map correlation were related differently to the two behaviors. We reversed the signs of reward decoding error and first long bout time so that higher values indicated better decoding and an earlier first long bout, and standardized each variable across the session pairs. For behavior *b*, either first long bout time (*T*) or exploration efficiency (*E*), we compared the two correlations,

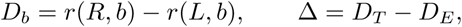

where *R* is reward decoding and *L* is level map correlation. *D_b_* asks which neural measure is more closely related to behavior *b*, and Δ asks whether that difference changes between the two behaviors.

To test *D_b_*= 0, we listed each session pair twice, once with *R* and once with *L* as the predictor *x*, each time with the same behavior value *y*, and fitted a model in which the two neural measures shared one slope,

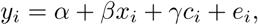

where *c_i_* is 1 for the copy using *L* and 0 otherwise. If *D_b_* = 0, this shared-slope model is complete. For each animal *a*, we then summed over its rows

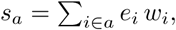

where *e_i_* is the residual of the shared-slope model and *w_i_* is the residual of *x_i_c_i_*, the term that would give *L* its own slope, after regression on the same terms. The test of Δ = 0 used the same construction with both behaviors listed together and the term that would let *D_b_* differ between behaviors. If the null hypothesis holds, each animal’s *s_a_* is equally likely to be positive or negative. We therefore compared the observed total *S*_score_ = Σ*_a_s_a_* with the totals Σ*_a_ε_a_s_a_* obtained for every choice of signs *ε_a_* = ±1 (all 2^14^ = 16,384 choices for the 14 animals).

### Statistics and reproducibility

Analyses were performed in Python 3.10.9 using NumPy 1.26.4, pandas 2.3.3, SciPy 1.10.0 and statsmodels 0.13.5. Statistical plots were generated with Matplotlib 3.7.0.

Unless stated otherwise, linear models were fitted by ordinary least squares, so every observation contributed equally to the fitted coefficients. Observations from the same animal can be correlated, so we treated each animal, rather than each observation, as the independent unit when estimating uncertainty. For a model 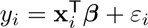 with fitted residuals *ε*^*_i_*, we first summed the contributions of each animal *g*,

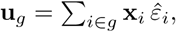

and then estimated the covariance of the coefficients from these *G* animal sums,

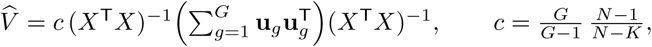

where *X* is the design matrix, *N* is the number of observations, *K* is the number of coefficients and *c* adjusts for the finite number of animals. Grouping these residual contributions by animal allowed repeated observations from the same animal to contribute jointly to uncertainty, without changing the fitted coefficients or observation weights. Each coefficient was divided by its standard error, the square root of the corresponding diagonal entry of 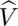, and compared with a Student’s *t* distribution with *G −* 1 degrees of freedom, where *G* was the number of animals.

Paired comparisons tested whether the mean within-pair difference differed from zero. When an animal contributed multiple pairs, these differences were clustered by animal. Bootstrap confidence intervals were obtained by sampling animals with replacement while keeping all observations from each sampled animal together. Percentile intervals were used unless stated otherwise. Tests were two-sided unless a directional test was specified.

No statistical method was used to predetermine sample sizes. Exclusions of sessions and session pairs from individual analyses, and the reason for each, are listed in Supplementary Table 1. Some comparisons used tests other than the clustered regression. Exact sign-flip tests summarized each animal by one score and compared the observed total with the totals obtained by flipping the signs of the animal scores in every possible combination. Shuffle tests repeated the analysis 10,000 times with neuron identities permuted within each session pair, and the one-sided *p* value was (1 + *k*)/10,001, where *k* was the number of shuffles whose statistic was at least as large as the observed value. Analyses with fewer shuffles did not derive *p* values from the shuffle distribution. Pearson correlations are reported as effect sizes; unless stated otherwise, their *p* values come from clustered regressions of the same variables.

When related tests formed a correction family, their *p* values were corrected with the Benjamini–Hochberg procedure, which controls the false discovery rate, the expected fraction of false discoveries among rejected hypotheses. For a family of *M* tests with raw *p* values ordered as *p*_(1)_ *≤ · · · ≤ p*_(_*_M_*_)_, the corrected values were

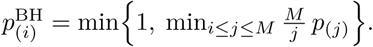

Taking the minimum over the same and larger ranks kept the corrected values in the order of the raw values, so several tests in one family could share one corrected value.

Figure asterisks denote *p <* 0.05 (*), *p <* 0.01 (**), and *p <* 0.001 (***). They represent corrected *p* values for family-controlled tests and raw *p* values for standalone tests.

## Acknowledgements

We thank Walter Gonzalez for help setting up the custom miniscope technique; Matthew Rosenberg for help with maze construction; Nicole Schweers for animal support and Joe Wekselblatt for breeding advice; Daniel Wagenaar of the Caltech Neurotechnology Lab for engineering assistance; the Caltech HPC for computing resources; Thanos Siapas for critical feedback; and all Meister lab and Lois lab members for their input. Generative AI tools (Claude, ChatGPT) were used for text editing, code generation, and debugging. The authors verified the final text and code and remain fully responsible for this work. This work was supported by grants from the Simons Collaboration on Global Brain (543015 to M.M.) and the National Institutes of Health (NIH) (R01NS111477, DP1NS155489 to M.M. and R01NS121918 to C.L.). The content is solely the responsibility of the authors and does not necessarily represent the official views of the NIH.

## Author contributions

J.W., M.M. and C.L. conceived the study. J.W. conducted the experiments and analyzed the data. J.W. wrote the paper with input from M.M. and C.L. M.M. and C.L. supervised all aspects of the work.

## Data and code availability

Source Data for all figures and tables, together with the code that rebuilds every figure, table and statistical result from them, are available at https://github.com/iamjiangwu/maze-lifelong-memory.

---

**Extended Data Figure 1.**
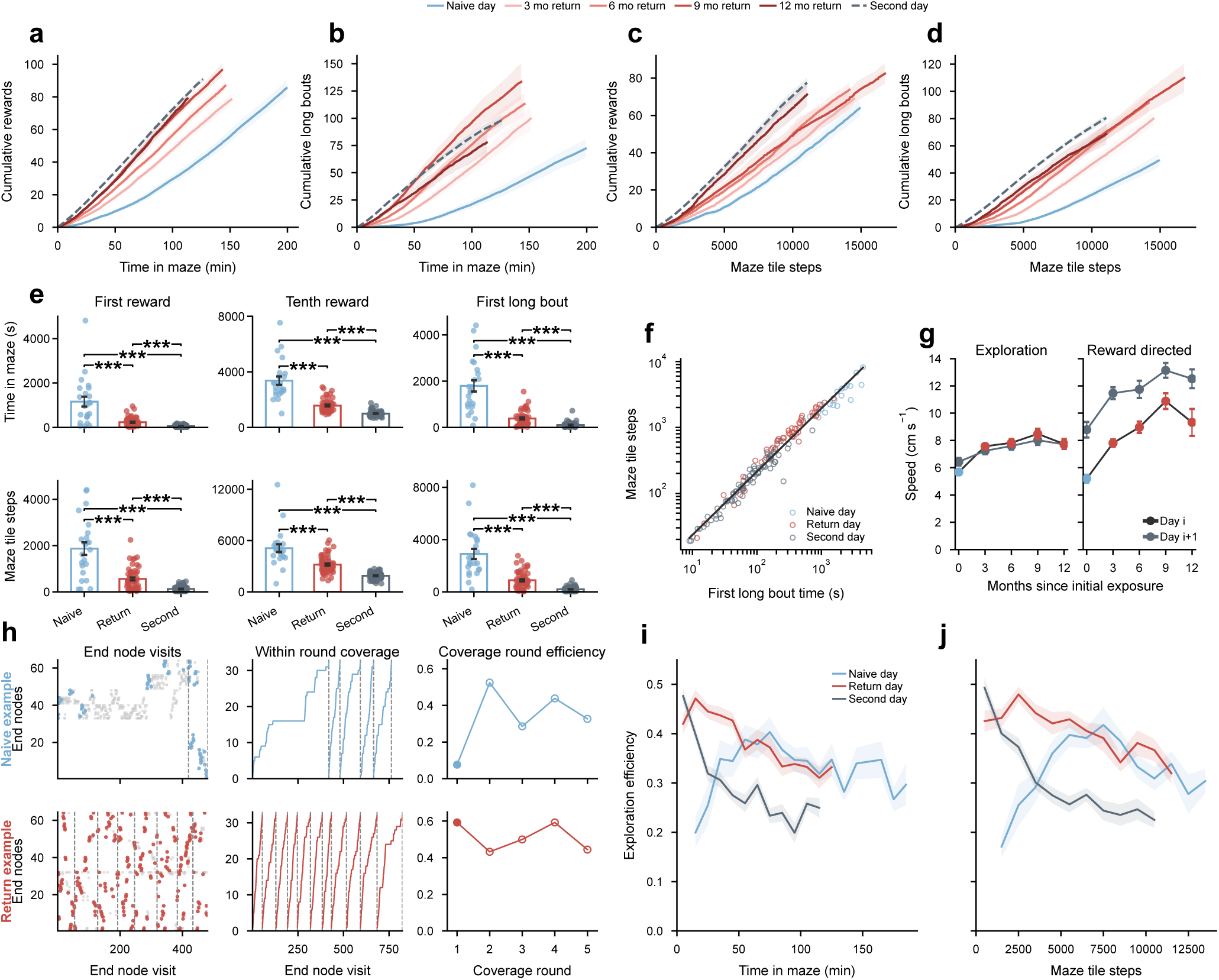
Additional reward and exploration measures for lifelong spatial memory. **a,** Cumulative rewards as a function of time in maze on the naive days, second days and return days after 3, 6, 9 and 12 months. Curves show session means ± s.e.m. through the last time point with at least 80% of sessions in each group (23, 74, 17, 17, 12 and 10 sessions, respectively). **b,** Cumulative long bouts over time in maze for the same groups as in **a**. **c,** Cumulative rewards as a function of maze tile steps. Curves show means ± s.e.m. over the tile-step range with at least 80% of sessions as in **a**. **d,** Cumulative long bouts as a function of maze tile steps for the same groups as in **a**. **e,** Columns show the first reward, tenth reward and first long bout; rows show elapsed time in maze in seconds (top) and maze tile steps (bottom). Points are sessions and bars show mean ± s.e.m. for naive day, return day and second day sessions (23, 56 and 74 sessions, respectively). Outcomes were analyzed after log(*x*) transformation using linear models with standard errors clustered by animal; all displayed comparisons were significant after BH correction (maximum corrected *p* = 1.7 × 10^−4^). **f,** Maze tile steps plotted against time in maze before the first long bout. Each point represents one session; both axes are logarithmic, and the black line shows the fitted relation. The slope was near unity (slope, 0.960; 95% CI, 0.93 to 0.98; 153 sessions from 29 animals). Of these animals, 23 contributed naive day sessions. The other six had initial sessions that were latently learned, so they contributed only later return day and second day sessions. Elapsed time and traversed distance therefore accumulated approximately proportionally across sessions. **g,** Nose speed during exploration (left) and reward-directed behavior (right) within the first 30 min of each session. The horizontal axis is months since initial exposure, and speed is measured in cm s^−1^. Symbols and error bars show mean ± s.e.m. on the first and second days of each visit, at 0, 3, 6, 9 and 12 months. For exploration, the respective sample sizes were 23, 17, 17, 12 and 10 Day *i* sessions and 18, 17, 17, 12 and 10 Day *i* + 1 sessions. For reward-directed behavior, they were 20, 17, 17, 12 and 10 sessions on Day *i* and 18, 17, 17, 12 and 10 sessions on Day *i* + 1. **h,** Repeated coverage rounds in example naive and return sessions. Left, successive end node visits; middle, the number of distinct end nodes visited within each round; right, exploration efficiency for each round. A round ends after 32 distinct end nodes have been visited, and its efficiency is 32 divided by the number of visits required. Dashed lines mark round boundaries. Filled symbols mark the first-round efficiency used in the main text. **i,** Exploration efficiency as a function of time in maze for naive day, return day and second day sessions. Curves show means ± s.e.m. in 10 min bins, ending at the last bin with at least 80% recording opportunity in each group. **j,** Exploration efficiency as a function of maze tile steps, summarized as in **i** in bins of 1,000 steps.

**Extended Data Figure 2.**
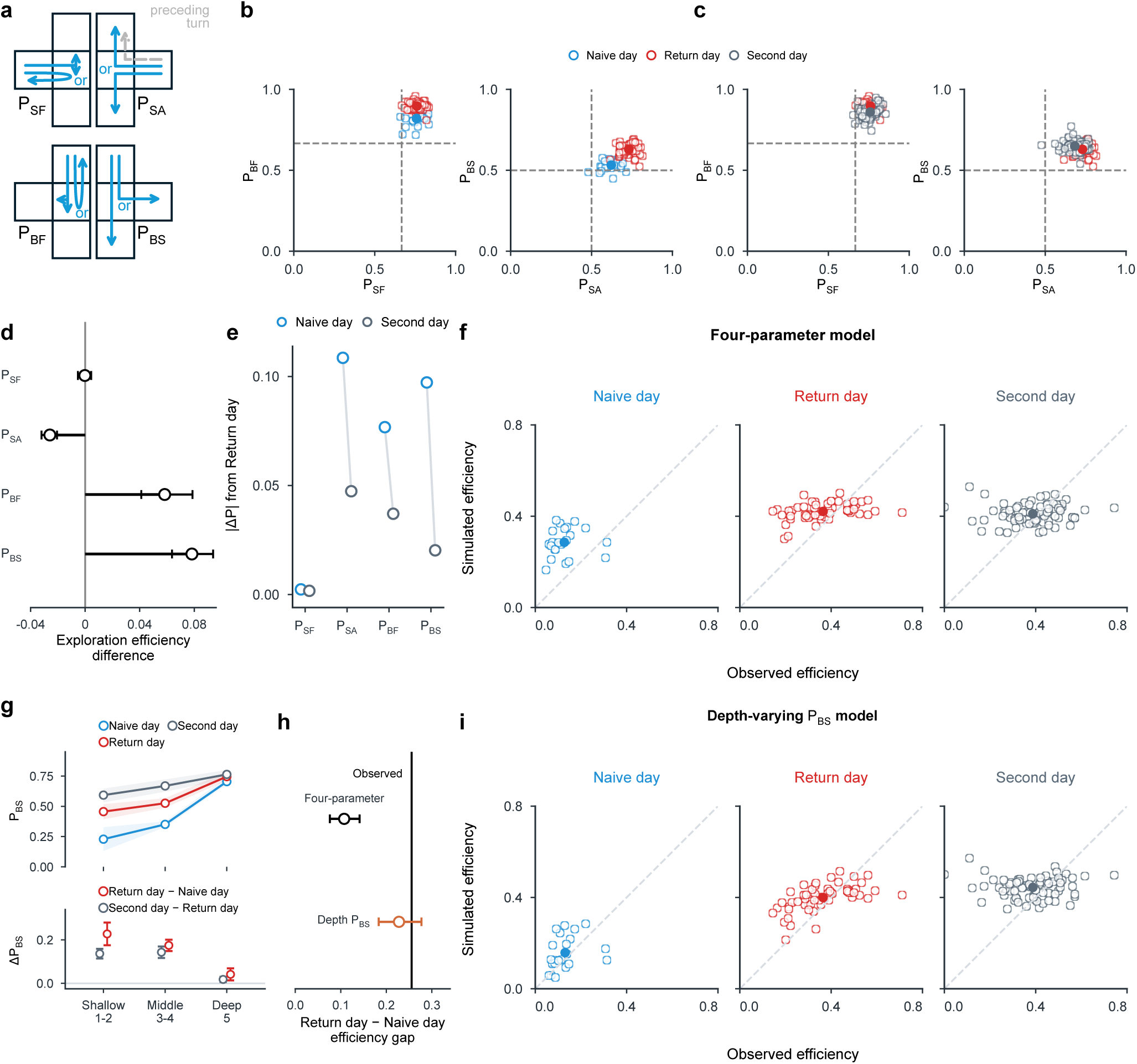
Junction turning biases and biased walk models of exploration efficiency. **a,** Four turning biases at junctions. P_SF_ is the probability of continuing from the stem into a branch rather than reversing. Conditional on continuing, P_SA_ is the probability of choosing the alternating branch. P_BF_ is the probability of leaving a previously sampled subtree rather than reentering it. Conditional on leaving, P_BS_ is the probability of continuing towards the stem rather than entering the sibling branch. **b,** Turning biases estimated for each session on naive and return days, restricted to sessions in which all four biases were estimable (23 sessions from 23 animals and 58 sessions from 18 animals, respectively). Hollow symbols are sessions, solid symbols are group means and dashed lines are equal choice references. Return minus naive differences were +0.002 for P_SF_ (BH corrected *p* = 0.84) and +0.08 for P_BF_ (corrected *p* = 4.9 × 10^−9^). The corresponding differences were +0.11 for P_SA_ (corrected *p* = 1.4 × 10^−8^) and +0.10 for P_BS_ (corrected *p* = 9.6 × 10^−11^). Biases were estimated from exploration choices during the first 30 min in the maze. In **b** and **c**, each session contributes one point to each plot. The left plots show P_SF_ on the horizontal axis and P_BF_ on the vertical axis; the right plots show P_SA_ and P_BS_ on those axes, respectively. Dashed lines mark equal-choice probabilities of 2/3 for the two forward choices among three exits and 1/2 for a conditional choice between two exits. **c,** Return day and second day biases, restricted to sessions in which all four biases were estimable (58 sessions from 18 animals and 74 sessions from 24 animals). Differences between second day and return day were −0.002 for P_SF_ (BH corrected *p* = 0.79) and −0.04 for P_BF_ (corrected *p* = 5.3 × 10^−6^). The corresponding differences were −0.05 for P_SA_ (corrected *p* = 6.5 × 10^−5^) and +0.02 for P_BS_ (corrected *p* = 0.0018). **d,** Contribution of each return day turning bias to simulated exploration efficiency. One bias at a time was replaced with the same animal’s naive value while the others were held fixed. The horizontal axis shows efficiency before replacement minus efficiency after replacement; positive values indicate that the observed return day bias improved exploration. Circles show estimates and horizontal bars show 95% confidence intervals (58 return sessions from 18 animals). Contributions were −0.0002 for P_SF_ (95% CI, −0.005 to +0.004), −0.026 for P_SA_ (−0.032 to −0.021), +0.058 for P_BF_ (+0.041 to +0.079), and +0.078 for P_BS_ (+0.064 to +0.094). **e,** For each turning bias, connected markers show the absolute differences of the naive day and second day means from the return day mean. Second day values were closer to return day values for all four biases. **f,** Observed exploration efficiency and efficiency simulated by the four-parameter biased walk. Hollow circles represent sessions and filled circles show group means; dashed diagonals mark equality (23 naive day, 58 return day and 74 second day sessions). Pearson *r* and mean bias were descriptive measures of session-level agreement (naive day, *r* = 0.09, bias = +0.16; return day, *r* = 0.38, bias = +0.06; second day, *r* = 0.05, bias = +0.02). **g,** Stemward exit probability P_BS_ across shallow, middle and deep non-root decision junctions. These included all junctions at the same tree depths corresponding to reward route decision nodes 1–2, 3–4 and 5, respectively, in Fig. 1d. The analysis used 22 naive day sessions from 22 animals, 58 return day sessions from 18 animals and 74 second day sessions from 24 animals with sufficient samples at all three depth groups. Upper curves show session means with interquartile bands; lower estimates show group contrasts with 95% CIs. BH corrected *p* values for return minus naive differences were 1.7 × 10^−9^, 1.7 × 10^−13^ and 0.0046 from shallow to deep. The corresponding corrected *p* values for differences between second day and return day were 2.0 × 10^−11^, 1.2 × 10^−10^ and 0.0045. **h,** Reconstruction of the observed difference of return and naive exploration efficiency using matched naive and return sessions from the same animals. The four-parameter biased walk reconstructed 42.0% of the observed gap (95% CI, 29.7 to 55.2%), whereas allowing P_BS_ to vary with depth reconstructed 89.1% (71.7 to 108.5%). The common four-parameter and depth-varying cohort contained 54 return sessions paired with 17 naive day sessions from 17 animals. Circles and horizontal intervals show reconstructed gaps and 95% confidence intervals; the vertical black line marks the observed return minus naive gap. **i,** Observed and simulated exploration efficiency for the model in which P_BS_ varied with depth. Hollow circles represent sessions and filled circles show group means; dashed diagonals mark equality. Included sessions had sufficient samples at all three depth groups, as in **g**. Pearson *r* and mean bias were *r* = 0.30 and +0.027 for naive day, *r* = 0.59 and +0.035 for return day, and *r* = 0.01 and +0.057 for second day.

**Extended Data Figure 3.**
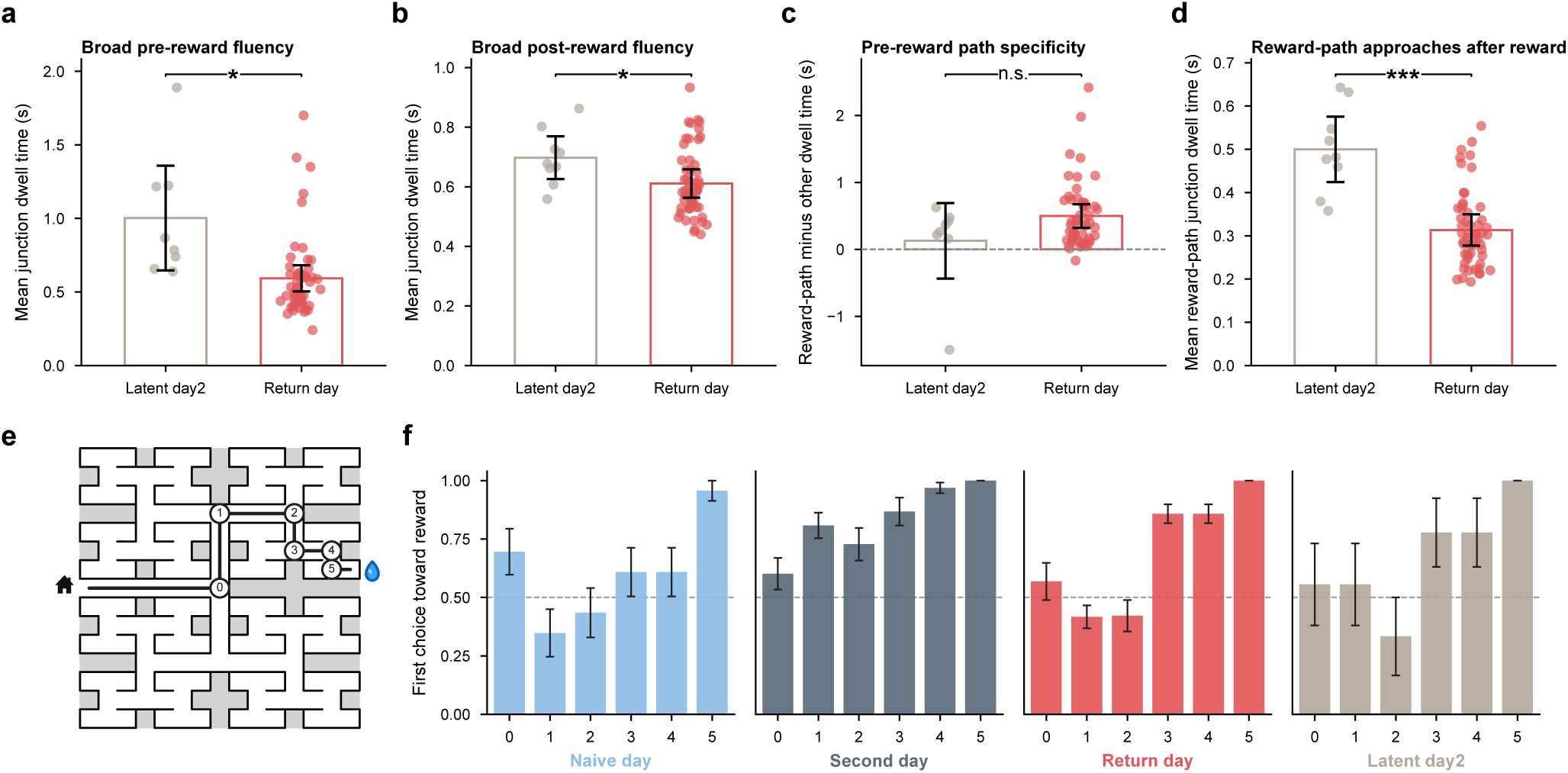
Reward path fluency and first visit route choices. **a,** Mean dwell time at internal junctions during exploration before the first reward encounter in latent day2 and return sessions. Return animals showed shorter junction dwell before reward, consistent with the hypothesis that animals were motivated for an anticipated reward before the reward was found (*β*_return_*_−_*_latent_ = −0.41 s; *β* stands for estimated difference; 55 return day and 8 latent day2 sessions from 20 animals with enough pre-reward samples; BH corrected *p* = 0.026). **b,** Mean exploratory junction dwell after the first reward. Return animals also showed shorter exploratory junction dwell after the first reward (*β*_return_*_−_*_latent_ = −0.09 s; 56 return day and 9 latent day2 sessions from 21 animals; corrected *p* = 0.032). **c,** Difference between dwell at junctions on the direct reward path and other junctions before the first reward. There was no supported difference specific to the reward path before reward (*β*_return_*_−_*_latent_ = +0.37 s; 53 return day and 8 latent day2 sessions from 20 animals; corrected *p* = 0.15). **d,** Junction dwell along the direct home to water path during reward-directed approaches after the first reward. Return animals traversed these junctions faster than latent day2 controls (*β*_return_*_−_*_latent_ = −0.19 s; 56 return day and 9 latent day2 sessions from 21 animals; corrected *p* = 2.2 × 10^−4^). In **a–d**, points are sessions, bars are group means and black error bars are 95% confidence intervals from regressions with standard errors clustered by animal. **e,** Six decision nodes used for the first visit reward path analysis. The direct home to water path follows the sequence left, right, right, left, right and left; circles numbered 0 to 5 mark the scored junctions. **f,** Probability that the first choice before reward at each junction was directed toward reward. The horizontal axis lists the six decision nodes numbered in **e**. Bars show means across animals ± s.e.m. for naive day, second day, return day and latent day2 sessions; the dashed line marks chance probability 0.5. Sample sizes were 23 sessions from 23 animals, 74 sessions from 24 animals, 56 sessions from 17 animals and 9 sessions from 9 animals, respectively. Second day animals favored the reward branch across most of the route. Return animals instead resembled latent day2 controls, including at choices favored by alternation; the home-to-water sequence (L R R L R L) contains four alternations in five turn transitions.

**Extended Data Figure 4.**
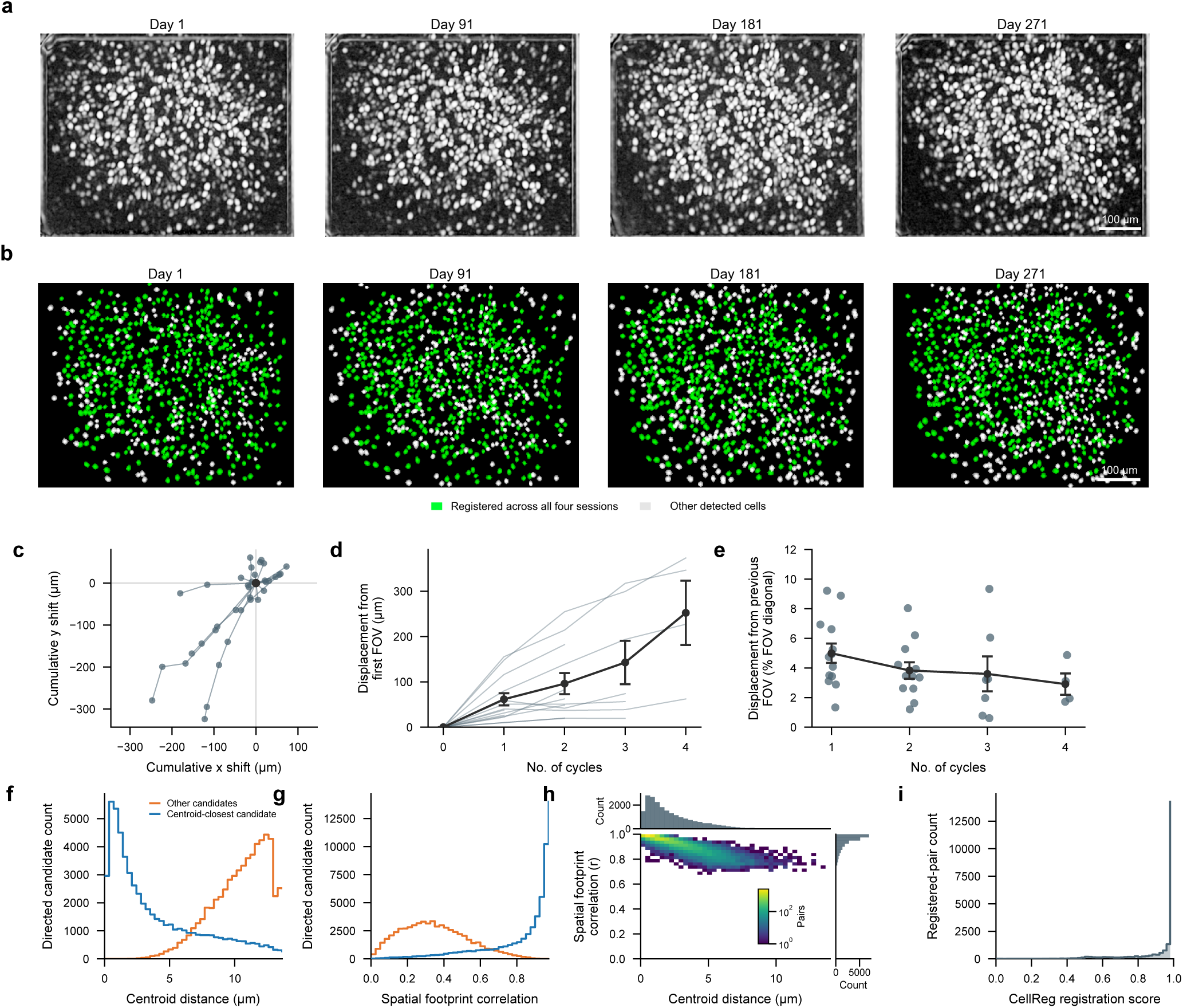
Cross-session field of view stability and neuron matching quality. **a,** Example longitudinal CA1 local correlation images on Day 1, Day 91, Day 181 and Day 271, showing a stable field of view across repeated long gaps. Scale bar, 100 *µ*m. **b,** The same fields of view with cells registered across all four cycles shown in green and other detected cells shown in white. **c,** Cumulative two dimensional field of view displacement across successive recording cycles. Each trajectory shows one animal, with the first field of view set to the origin (14 animals). **d,** Displacement from the first field of view across recording cycles. Thin lines show individual animals and black points show mean ± s.e.m. Mean displacement at cycle counts 1 to 4 was 61.6, 96.1, 142.8 and 252.3 *µ*m, respectively (*n* = 13, 12, 7 and 4 animals). **e,** Displacement from the preceding field of view across recording cycles, expressed as a percentage of the preceding field of view diagonal. Points denote animals; black symbols and the connecting line show means across animals at each cycle, with s.e.m. error bars. Mean displacement at cycle counts 1 to 4 was 4.99%, 3.83%, 3.60% and 2.91%, respectively (*n* = 13, 12, 7 and 4 animals; pooled median, 3.45%). **f,** Centroid distance distributions for candidate neuron matches within the 14 *µ*m search radius, evaluated in both directions within each session pair. The nearest candidate by centroid was classified as centroid closest and all remaining candidates as other candidates. The analysis comprised 57,754 centroid closest and 61,690 other candidate observations from 36 session pairs and 14 animals; median distances were 2.62 and 10.93 *µ*m, respectively. **g,** Spatial footprint correlations for the same directed candidates in **f**. Median correlations were 0.930 for centroid closest candidates and 0.333 for other candidates. **h,** Joint distribution of centroid distance and spatial footprint correlation after alignment for 21,412 neuron pairs registered with CellReg. Color denotes logarithmically scaled raw bin count and the marginal histograms show linear raw counts. Median centroid distance was 1.68 *µ*m and median footprint correlation was 0.959. **i,** Distribution of CellReg registration scores for the same registered neuron pairs. The median was 0.997 and 74.7% were at least 0.95.

**Extended Data Figure 5.**
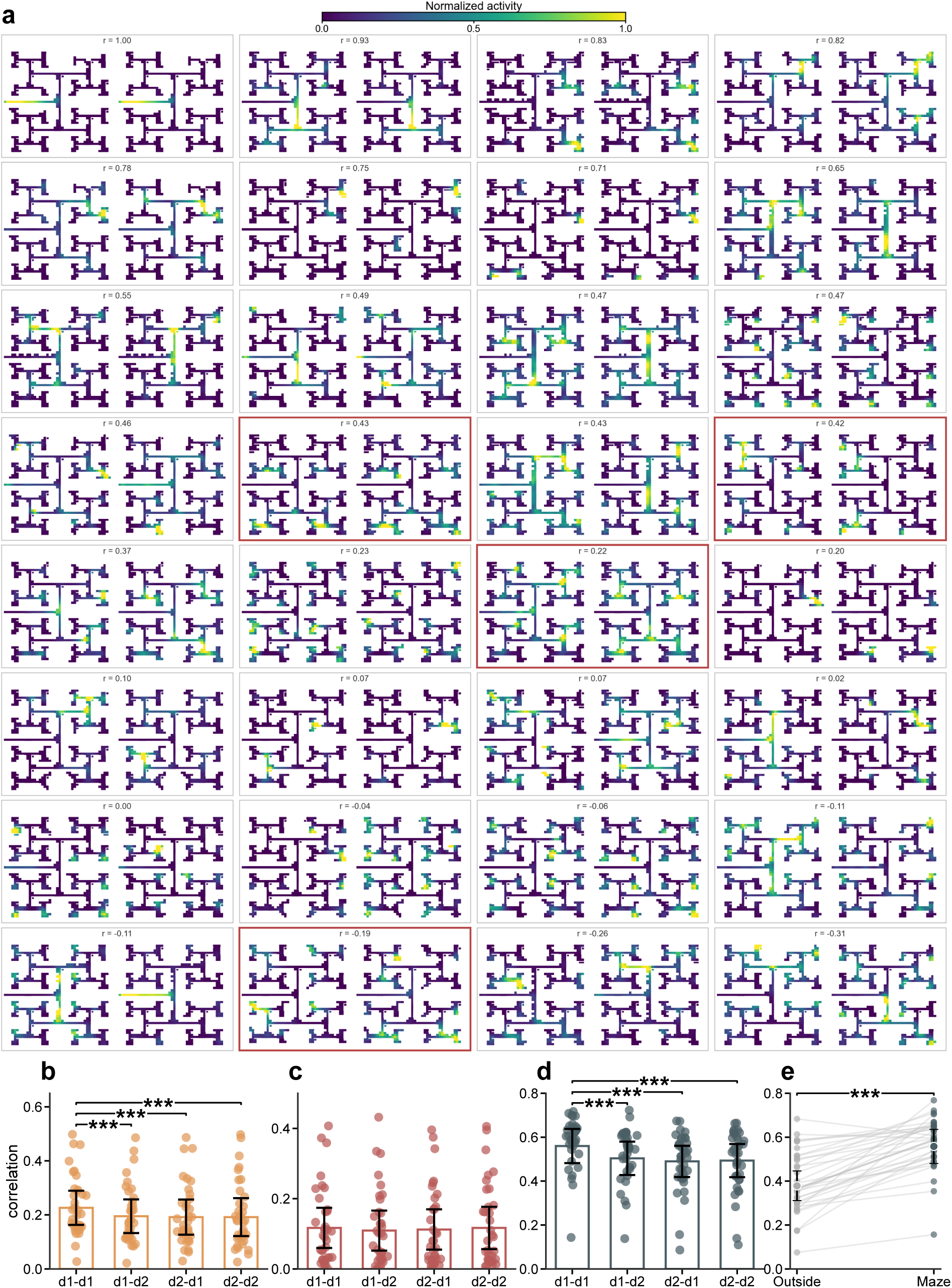
Examples and controls for long gap neural stability. **a,** Thirty-two illustrative place map pairs from neurons matched across 3 month gaps, ordered from higher to lower correlation from left to right and top to bottom. Each pair shows the earlier map on the left and the later map on the right. Each map was independently normalized to its 99th percentile activity for display; color denotes normalized activity, and the Pearson correlation is shown above each pair. Red outlines identify the four examples shown in Fig. 2b. **b,** Long gap PV correlations across four day combinations. The labels d1–d1, d1–d2, d2–d1 and d2–d2 denote comparisons of Day *i* with Day *i* + 90, Day *i* with Day *i* + 91, Day *i* + 1 with Day *i* + 90, and Day *i* + 1 with Day *i* + 91, respectively. Points show session pairs, bars show means across session pairs and error bars show 95% confidence intervals using standard errors clustered by animal (36 session pairs from 14 animals). PV correlations were highest for d1–d1 comparisons (means, 0.23, 0.20, 0.19 and 0.19). d1–d1 exceeded d1–d2, d2–d1 and d2–d2 (BH corrected *p* values were 3.7 × 10^−4^, 4.9 × 10^−4^ and 3.7 × 10^−4^, respectively). The other three pairwise comparisons were not significant. **c,** Same as **b** but for place map correlation. Place map correlation means were 0.12, 0.11, 0.11 and 0.12 across the four day combinations, respectively. No significant differences were detected between any comparisons. **d,** Same as **b** but for ensemble rate correlation. Ensemble rate correlations were highest for d1–d1 comparisons (means, 0.56, 0.50, 0.49 and 0.49). d1–d1 exceeded d1–d2, d2–d1 and d2–d2 (BH corrected *p* values were 7.3 × 10^−4^, 7.3 × 10^−4^ and 7.3 × 10^−4^, respectively). The other three pairwise comparisons were not significant. **e,** Ensemble rate correlations outside the maze and during maze behavior with equal samples for d1–d1 long gap pairs. Samples from outside the maze include the home cage and connecting tunnel. Gray lines connect the same session pair observations; black points show means across pairs and error bars show 95% confidence intervals from regressions with standard errors clustered by animal. Correlation was 0.38 outside the maze (95% CI, 0.31 to 0.45) and 0.56 inside the maze (0.48 to 0.64). The paired difference was 0.18 (95% CI, 0.14 to 0.22; 36 pairs from 14 animals; *p* = 5.1 × 10^−7^).

**Extended Data Figure 6.**
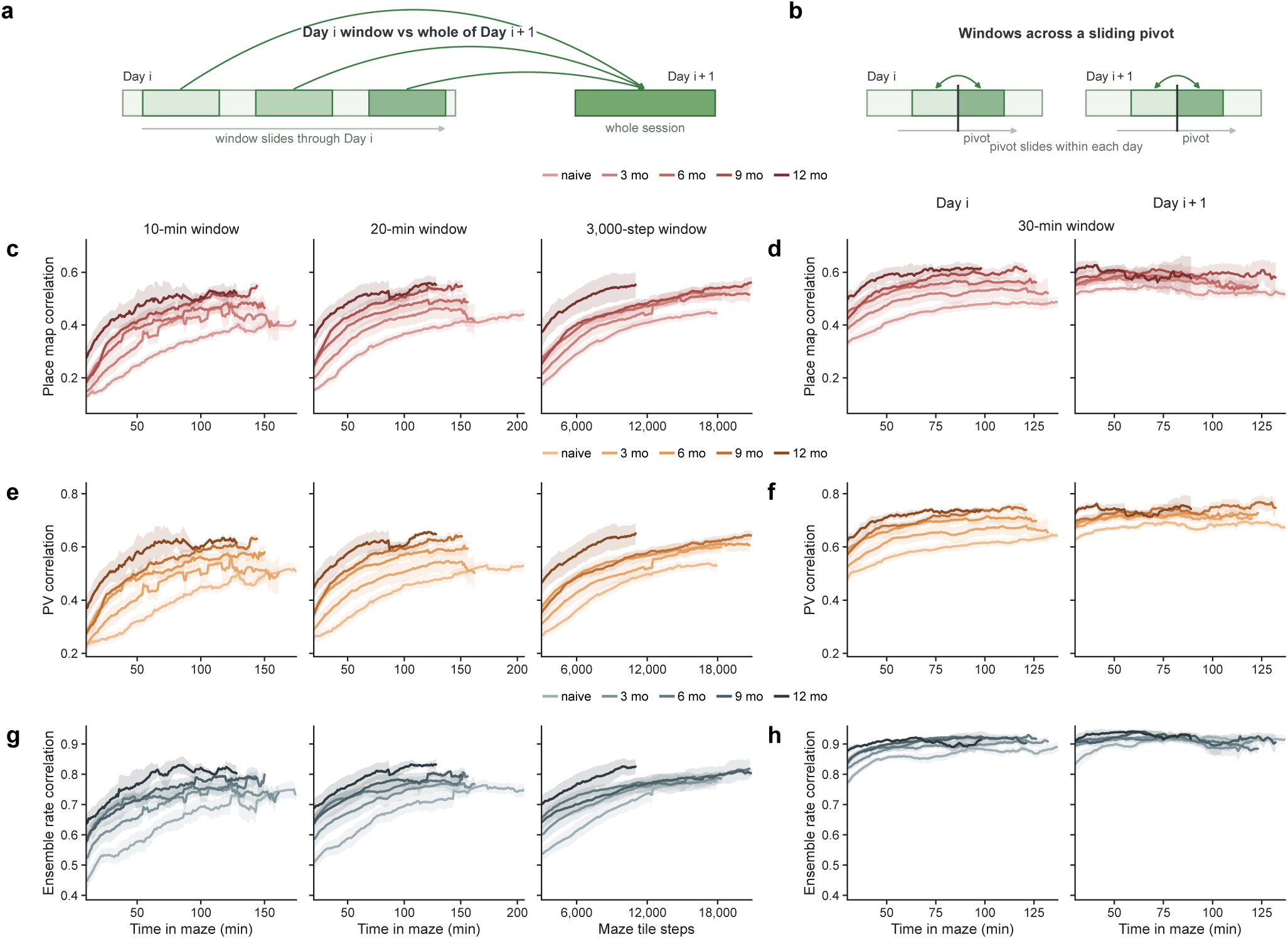
Convergence within recording cycles and local stability within sessions. **a,** Schematic of the fixed reference analysis. Moving windows from the Day *i* session were compared with the full Day *i* + 1 representation from the same recording cycle. **b,** Schematic of the sliding pivot analysis. Two abutting windows before and after a pivot were compared as the pivot moved through Day *i* or Day *i* + 1. **c,** Place map correlation between Day *i* windows and the full Day *i* + 1 representation used as the reference, using windows of 10 min, 20 min or 3,000 tile steps. **d,** Place map correlation between the 30 min windows before and after a sliding pivot. Whole-curve variability was lower on Day *i* + 1 than on Day *i* (within-curve s.d., 0.025 versus 0.047 *r*; BH corrected *p* = 2.2 × 10^−7^). Separate linear tests on Day *i* show a 0.11 *r* per hour increase (corrected *p* = 1.7 × 10^−9^), while Day *i* + 1 shows no significant increase (corrected *p* = 1.0). **e,** PV correlation between Day *i* windows and the full Day *i* + 1 representation used as the reference. **f,** PV correlation across the 30 min sliding pivot. Whole-curve variability was lower on Day *i* + 1 than on Day *i* (within-curve s.d., 0.027 versus 0.044 *r*; BH corrected *p* = 2.6 × 10^−5^). Separate linear tests on Day *i* show a 0.10 *r* per hour increase (corrected *p* = 1.7 × 10^−9^) and Day *i* + 1 shows a marginal increase of 0.03 *r* per hour (corrected *p* = 0.0054). **g,** Ensemble rate correlation between Day *i* windows and the full Day *i* + 1 representation used as the reference. **h,** Ensemble rate correlation across the 30 min sliding pivot. Whole-curve variability did not differ between Day *i* + 1 and Day *i* (within-curve s.d., 0.026 versus 0.031 *r*; BH corrected *p* = 0.16). Separate linear tests on Day *i* show a 0.05 *r* per hour increase (corrected *p* = 3.1 × 10^−8^), while Day *i* + 1 shows no significant increase (corrected *p* = 0.44). Across **c–h**, color opacity indicates cumulative experience. Curves show means across animals with s.e.m. for each experience group. Fixed reference and sliding pivot analyses both used 59 recording cycles from 22 animals.

**Extended Data Figure 7.**
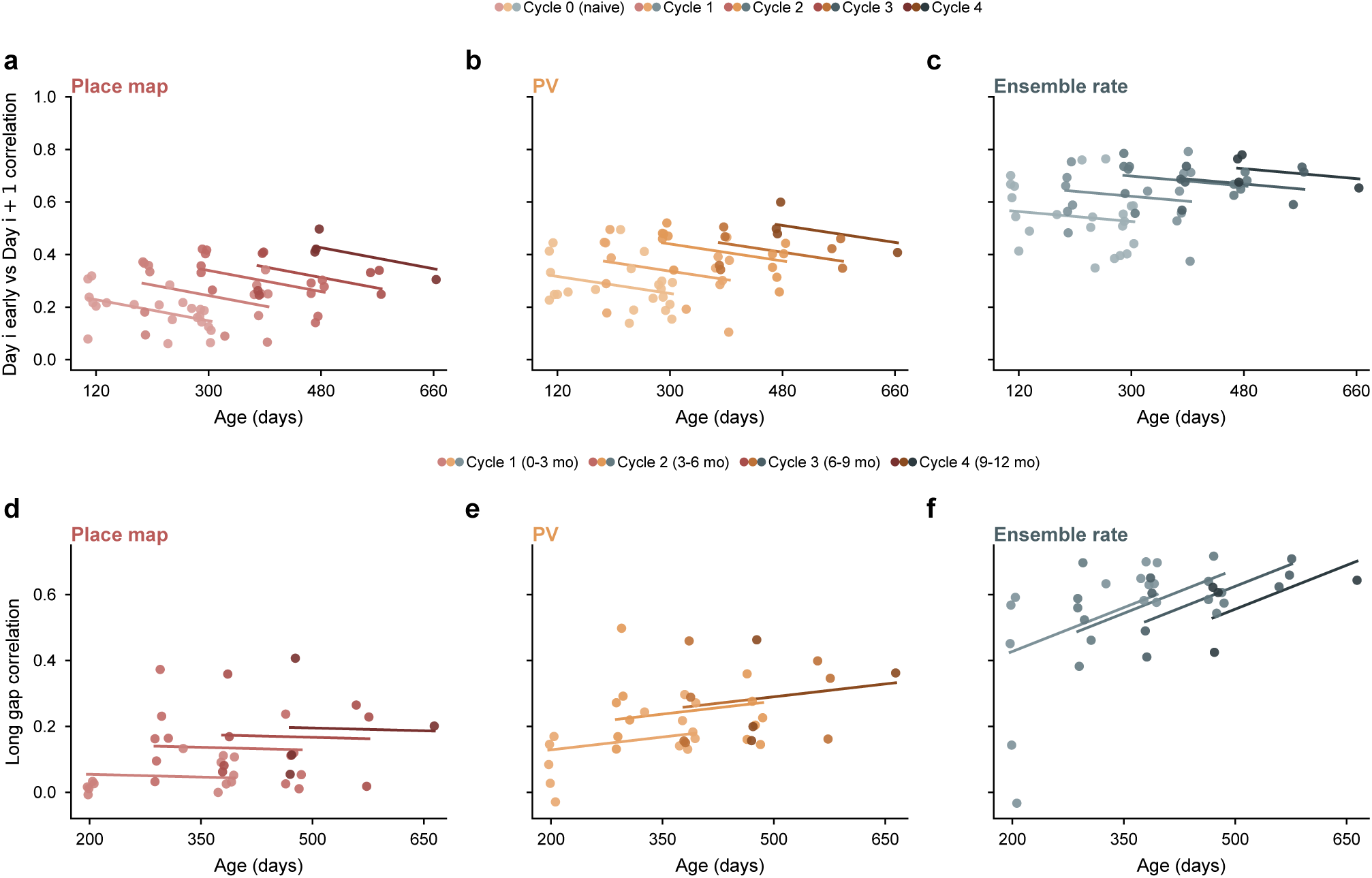
Age associations with map stability. **a–c,** Place map, PV and ensemble rate correlations between the first 30 min of Day *i* and the full Day *i* + 1 representation, plotted against age on Day *i*. The analysis included 59 observations from 22 animals, each representing one animal in one recording cycle. Solid circles show observations, colors indicate cycle stage, and lines show a common age slope with a separate intercept for each cycle stage. Per 30 d of age, place map correlation decreased by 0.013 (95% CI, −0.023 to −0.004; BH corrected *p* = 0.024). PV correlation decreased by 0.011 (−0.022 to 3.1 × 10^−5^; corrected *p* = 0.076). Ensemble rate correlation decreased by 0.006 (−0.017 to 0.004; corrected *p* = 0.24). Only the place map correlation decreased significantly. **d–f,** Long gap place map, PV and ensemble rate correlations plotted against age in the later session. The analysis included 36 pairs from 14 animals. Solid circles are session pairs, colors denote the later recording cycle, and lines show the age association adjusted for recording cycle transition. Per 30 d, the slopes were −0.002 for place maps (95% CI, −0.019 to 0.016; BH corrected *p* = 0.84) and +0.008 for PVs (−0.012 to 0.027; corrected *p* = 0.81). The ensemble rate slope was +0.027 (0.004 to 0.049; corrected *p* = 0.098). No association was significant after correction.

**Extended Data Figure 8.**
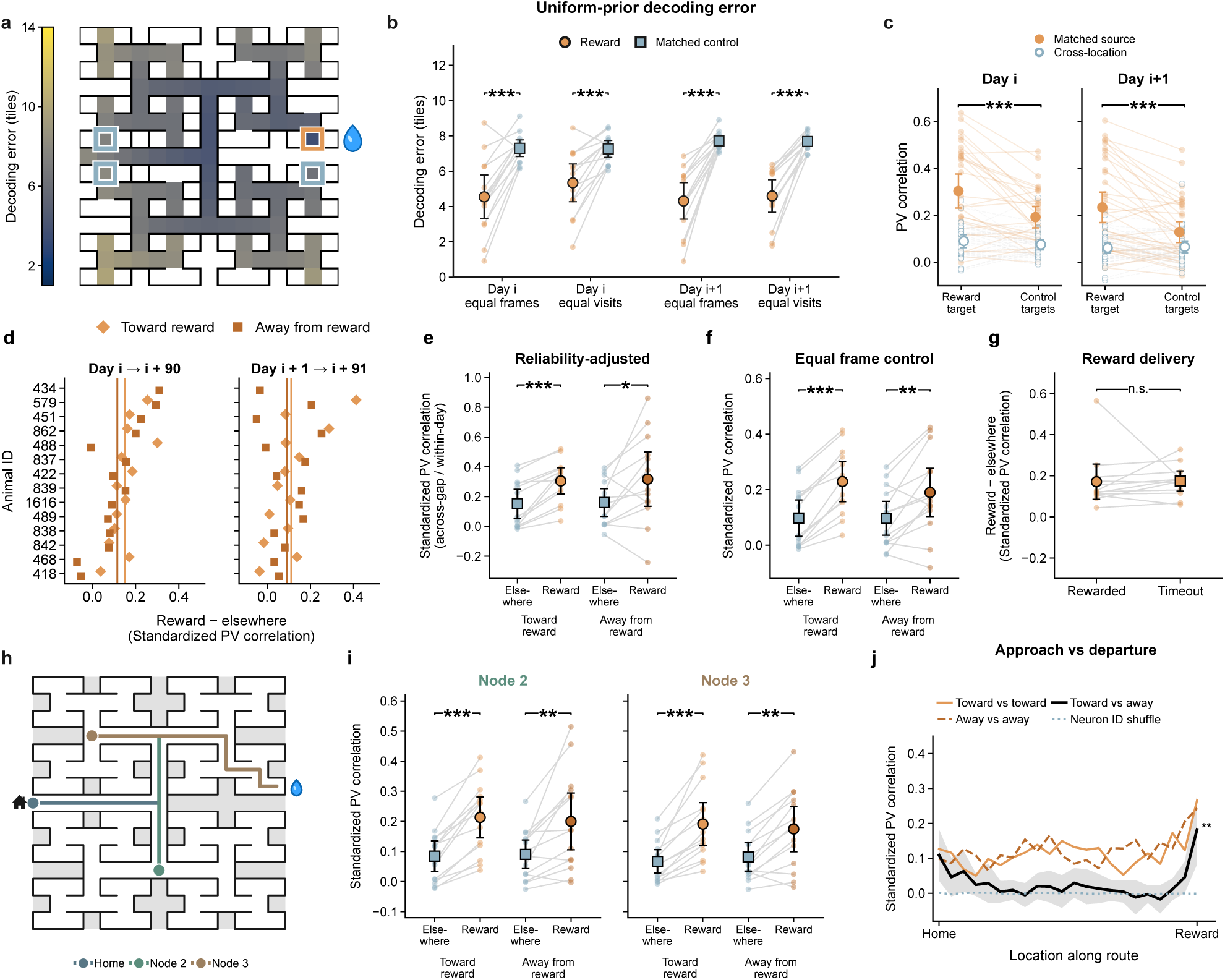
Reward code stability across sampling and route controls. **a,** Mean decoding error at each maze location, averaged equally across animals. Color shows decoding error in maze tiles. The map includes 104 maze tiles represented in all 14 animals and uses 30 pairs with rewarded Day *i* source sessions. The water drop marks the water port; the orange square outlines the reward decoder tile analyzed in **b**, and blue-gray squares outline the matched control tiles. **b,** Decoding error at the reward location and matched control locations after equalizing either recording frames or visits in both sessions, using a uniform spatial prior. The horizontal groups identify the source day and sampling control. Orange circles denote the reward location and blue-gray squares the control locations. With Day *i* sources, differences between control and reward tiles were 2.8 tiles with equal frames (95% CI, 1.8 to 3.7; 30 pairs from 14 animals; BH corrected *p* = 3.1 × 10^−4^). With equal visits, the difference was 1.9 tiles (1.1 to 2.8; corrected *p* = 3.1 × 10^−4^). With Day *i* + 1 sources, the corresponding differences were 3.4 tiles (2.5 to 4.4; 36 pairs from 14 animals; corrected *p* = 1.2 × 10^−4^) and 3.1 tiles (2.3 to 3.9; corrected *p* = 1.2 × 10^−4^). Differences were tested with one-sided exact sign-flip tests on animal means, with BH correction. Small points and connecting lines are animal means; large circles and squares with black error bars show means across animals and 95% confidence intervals. **c,** PV correlation at reward and control locations across a 3-month gap, using Day *i* (left) or Day *i* + 1 (right) as the earlier session. The horizontal axis identifies the later-session target location; the vertical axis is the correlation with the earlier-session source. Filled orange symbols compare each target with the same location in the earlier session. Open blue-gray symbols compare the reward target with the earlier control locations, or control targets with the earlier reward location. Small points represent session pairs, and lines connect the reward and control values from the same pair. Large symbols and error bars show means and descriptive 95% confidence intervals. Same location correlations were higher at reward than at controls by 0.11 for Day *i* (95% CI, 0.06–0.16; 30 pairs from 14 animals; BH-corrected *p* = 3.3 × 10^−4^) and 0.11 for Day *i* + 1 (0.06–0.15; 36 pairs from 14 animals; corrected *p* = 3.3 × 10^−4^). Tests and intervals for the paired differences accounted for repeated pairs within animals. **d,** Port advantage in individual animals. The horizontal axis shows standardized PV correlation at the rewarded end node minus its mean elsewhere along the route; each row is one animal. Light-orange diamonds denote approach and dark-orange squares departure. The left plot compares Day *i* with Day *i* + 90 and the right plot compares Day *i* + 1 with Day *i* + 91. Vertical colored lines show the corresponding means across animals. The port advantage was positive in all 13 animals with enough approach samples and 11 of 14 animals with enough departure samples on the first day. On the second day, mean advantages were 0.11 and 0.09, respectively (28 and 27 session pairs; 13 and 14 animals; BH-corrected *p* = 0.0039 and 0.0045). **e,** Standardized PV correlation across the 90-day gap relative to within-day reliability, shown separately at the rewarded end node and elsewhere for each direction. Each session’s linear runs were separated into odd and even runs. Within-day similarity was measured by correlating the population vectors from these two sets in each session. Across-gap similarity was the average of all four combinations between the two sessions. After averaging within each animal, across-gap similarity was divided by the geometric mean of the two within-day similarities. The paired comparison used 13 animals in each direction. The differences between port and elsewhere on the route were 0.15 during approach (95% CI, 0.10–0.21; BH-corrected *p* = 4.9 × 10^−4^) and 0.16 during departure (0.02–0.30; corrected *p* = 0.029). In **e–g** and **i**, small connected points show each animal’s paired values; large symbols and error bars show means across animals and 95% confidence intervals. **f,** Standardized PV correlation at the rewarded end node and elsewhere after equalizing the number of retained frames across tiles within each session. The port advantage was 0.13 during approach (95% CI, 0.10–0.16; 13 animals; BH-corrected *p* = 4.9 × 10^−4^) and 0.09 during departure (0.04–0.15; 14 animals; corrected *p* = 0.0053). **g,** Port advantage on rewarded and timeout approaches. The vertical axis shows standardized PV correlation at the rewarded end node minus its mean elsewhere; lines connect the two values for each animal (25 pairs from 12 animals). Rewarded visits had water delivery within 3 s of entry into the rewarded end node; timeout visits did not. Their difference was −0.003 (rewarded minus timeout; 95% CI, −0.089 to 0.083; *p* = 0.95). **h,** The three routes analyzed in both directions, connecting the water port to home, node 2 or node 3. Colored paths identify the three routes; green and brown match the node 2 and node 3 headings in **i**. **i,** Standardized PV correlation at the rewarded end node and elsewhere on the node 2 route (left; 30 pairs from 14 animals) and node 3 route (right; 29 pairs from 14 animals), with approach and departure shown separately. The differences were 0.13 and 0.11 for approach and departure on the node 2 route (BH-corrected *p* = 4.9 × 10^−4^ and 0.0015), and 0.12 and 0.09 on the node 3 route (corrected *p* = 4.9 × 10^−4^ and 0.0014). **j,** Standardized PV correlation across the 90-day gap as a function of physical location, from home on the left to the port on the right. Solid orange and dashed dark-orange curves compare approach with approach and departure with departure, respectively. The black curve compares opposite directions across the gap, averaging approach on Day *i* with departure on Day *i* + 90 and the reverse pairing. All curves use the same matched neurons and session pairs (26 pairs from 13 animals). The dotted blue-gray curve is the neuron-ID shuffle for the cross-direction comparison. Shading around the black curve shows the 95% confidence interval across animals. Cross-direction correlation was 0.18 at the rewarded end node and 0.02 elsewhere (difference, 0.16; 95% CI, 0.07–0.26; *p* = 0.0032).

**Extended Data Figure 9.**
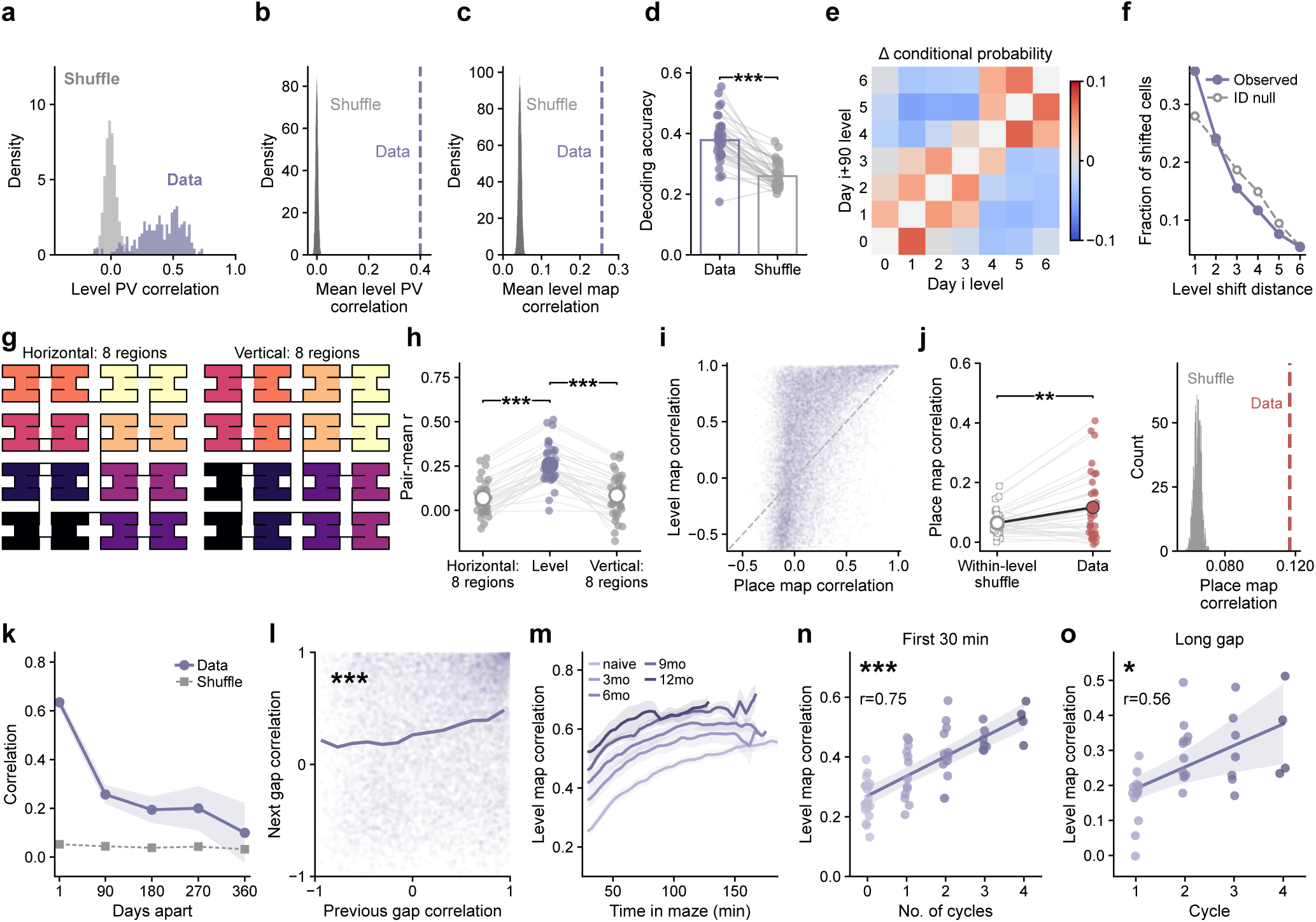
Additional analyses of the persistent level code. **a,** Distribution of level PV correlations across a 90-day gap. Each observation compares the activity of matched neurons at the same tree level in a session pair. Purple shows the data and gray the neuron ID shuffle; the vertical axis is probability density (248 level pair observations from 36 session pairs and 14 animals; mean correlation, 0.40). **b,** Mean level PV correlation compared with neuron ID shuffles. The gray histogram shows the probability distribution of mean correlations from 10,000 shuffles; the purple dashed line marks the observed mean across session pairs (0.40). **c,** Mean level map correlation compared with neuron ID shuffles. The gray histogram shows mean correlations from 10,000 shuffles and the purple dashed line marks the observed mean across session pairs (data, 0.26; one-sided permutation *p* = 1.0 × 10^−4^). **d,** Cross-session level decoding accuracy, measured as the fraction of target-session frames assigned to the correct tree level. The shuffle retained unequal occupancy across levels and the training prior rather than assigning equal probability to all seven levels. Points represent session pairs, and lines connect observed and neuron-ID-shuffled accuracy; (data, 0.38 ± 0.01; shuffle, 0.26 ± 0.01; *n* = 36 session pairs from 14 animals; one-sided regression with standard errors clustered by animal, *p* = 2.2 × 10^−6^). **e,** Preferred level transitions among the 13,679 matched neuron observations whose preferred level changed, from 18,409 observations across 36 session pairs and 14 animals. Columns identify the earlier preferred level and rows the later preferred level. Color shows observed minus neuron-ID shuffled transition probability. Shuffled probabilities were averaged over 10,000 permutations within each pair. Probabilities were normalized among neurons that changed level. **f,** Distance between earlier and later preferred levels for the same neurons as in **e**. The vertical axis gives the fraction of neurons that shifted by each distance. The solid curve shows the observed distribution; the dashed curve shows the permutation mean. Adjacent-level shifts exceeded the permutation expectation by 0.070 (95% CI, 0.054 to 0.086; BH corrected *p* = 2.6 × 10^−7^), and mean shifts were 0.21 levels shorter than expected (0.16 to 0.26; corrected *p* = 2.8 × 10^−7^). **g,** Horizontal and vertical partitions of the maze into eight regions. Colors identify maze tiles combined into each region. **h,** Long gap map correlations for the two spatial partitions in **g** and the level representation in Fig. 5a. Small connected points show session pairs and large points show means across pairs (36 pairs from 14 animals). Mean correlations were 0.07, 0.26 and 0.08 for horizontal, level and vertical maps, respectively. Level map correlations exceeded both spatial controls (one-sided regressions with standard errors clustered by animal; BH corrected *p* = 2.7 × 10^−8^ and *p* = 1.3 × 10^−6^, respectively). **i,** Place map correlation and level map correlation for the same matched neurons (18,409 observations from 36 session pairs and 14 animals). Darker regions contain more overlapping points. The dashed diagonal marks equality; points above it have higher level map than place map correlation. **j,** Place map correlation after shuffling locations within each tree level. Left, lines connect each session pair’s observed and shuffled correlations; large symbols show means across pairs. Right, the gray histogram counts 1,000 shuffled cohort means and the red dashed line marks the observed mean (difference, 0.05; 95% CI, 0.01 to 0.09; 36 pairs from 14 animals; one-sided regression with standard errors clustered by animal, *p* = 0.0072). **k,** Level map correlation as a function of the interval between sessions. Points and shading show means and 95% confidence intervals across session pairs; the dashed curve shows neuron ID shuffle means. The 1 day comparison is within a recording cycle (51 session pairs from 14 animals). Longer intervals compare Day *i* sessions across recording cycles (36, 22, 11 and 4 session pairs from 14, 11, 7 and 4 animals at 90, 180, 270 and 360 d, respectively). Comparisons longer than three months included intervening maze visits. **l,** Level map stability across consecutive gaps. Each faint point shows one neuron’s correlation across the previous gap and the next gap. The solid curve connects mean next-gap correlations within equal-width bins of previous-gap correlation (8,098 matched neuron observations from 22 three-session chains and 11 animals; regression slope, 0.20; BH corrected *p* = 8.1 × 10^−5^; standard errors clustered by animal). **m,** Level map correlation between moving 30 min windows on Day *i* and the full Day *i* + 1 session. The horizontal axis gives time in maze at the end of each window. Curves show smoothed means across animals and shaded bands show s.e.m.; darker colors indicate greater prior experience (59 recording cycles from 22 animals, with 22, 14, 12, 7 and 4 animals at naive, 3, 6, 9 and 12 month experience, respectively). **n,** Level map correlation between the first 30 min of Day *i* and the full Day *i* + 1 session, plotted against the number of completed recording cycles. Each point represents one animal in one cycle; zero denotes the naive visit (59 observations from 22 animals; Pearson *r* = 0.75; BH corrected *p* = 2.9 × 10^−10^). **o,** Level map correlation across successive recording cycles, plotted against the later cycle number (37 session pairs from 14 animals, including one short interval pair; Pearson *r* = 0.56; BH corrected *p* = 0.017). Each point represents one session pair. In **n** and **o**, lines are linear fits and shading shows 95% confidence intervals for the fitted means; tests and intervals account for repeated observations from each animal.

**Extended Data Figure 10.**
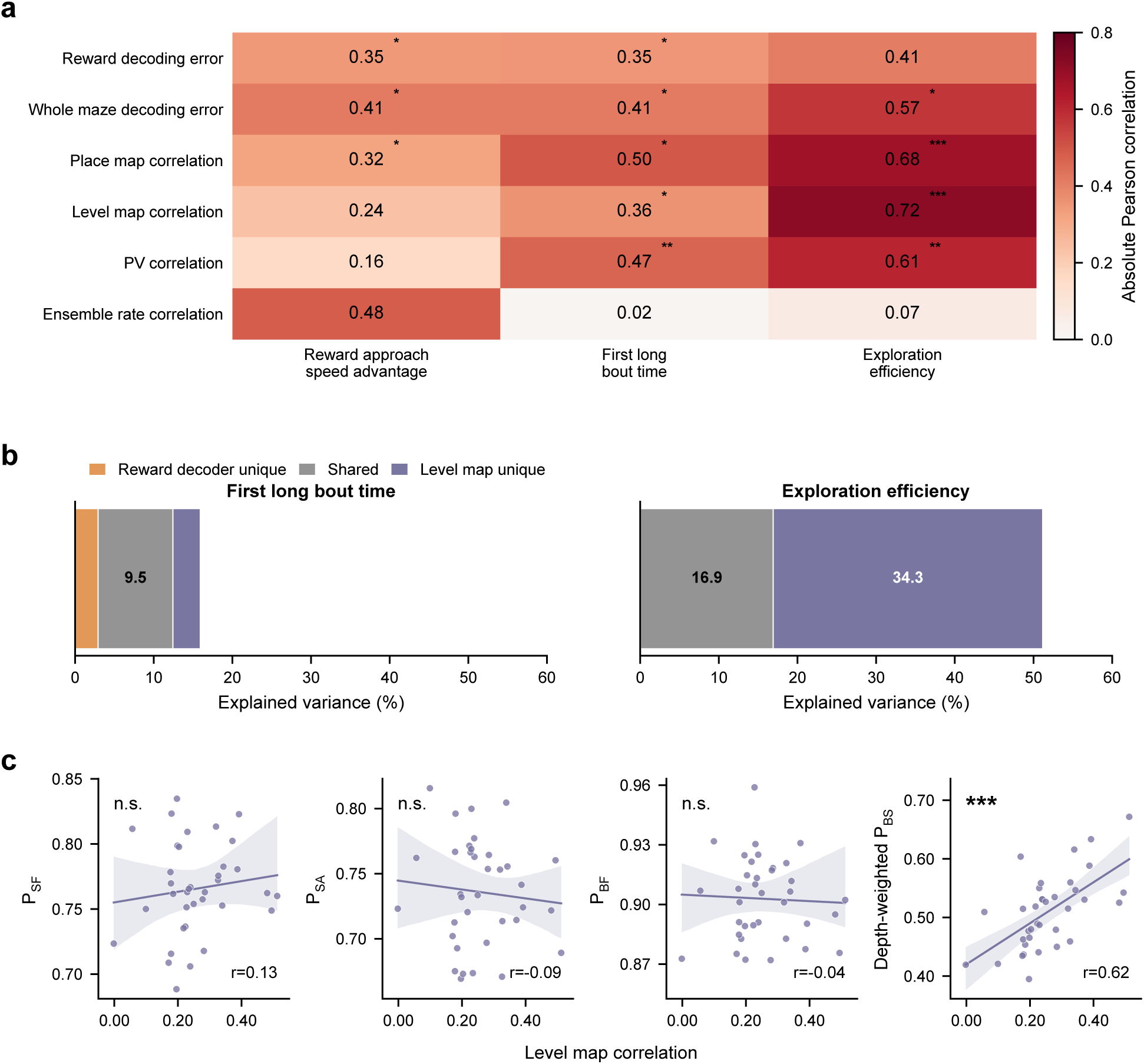
Neural preservation and later behavior. **a,** Strength of the associations between six neural readouts across a 3 month gap (rows) and three behaviors measured in the later session (columns). Each number and its cell color show the absolute Pearson correlation, *|r|*, across the same 30 session pairs from 14 animals. Reward approach speed advantage is the log ratio of median nose speed during approaches to the rewarded end node and to the three symmetrically placed unrewarded end nodes. All neural readouts used the same number of neurons from all session pairs, and both decoder rows used 2,000 frames per pair, sampled across the whole later session. Stars indicate BH corrected *p* values from regressions with standard errors clustered by animal. **b,** Commonality partition of variance jointly associated with reward decoding error and level map correlation. Bar segments show the percentage of behavioral variance associated uniquely with reward decoding (orange), uniquely with level map correlation (purple), or shared by the two (gray). The left plot concerns time to the first long bout and the right plot concerns exploration efficiency. Separate tests compared the strengths of the neural–behavior correlations. Across the same 30 session pairs from 14 animals, the relative strength of the two neural associations differed between behaviors (two-sided sign-flip score test using one score per animal, BH corrected *p* = 0.012). Reward decoding and level map correlation had similar associations with earlier long bouts (difference in *r*, −0.01 for reward decoding minus level map correlation; corrected *p* = 0.95). Level map correlation was more strongly associated with exploration efficiency than reward decoding was (difference in r, 0.30; corrected *p* = 0.044). **c,** Level map correlation across the gap against turning biases on return. From left to right, the plots show P_SF_, P_SA_, P_BF_ and depth-weighted P_BS_, defined in Extended Data Fig. 2a. Each point represents one session pair (36 pairs from 14 animals). Turning biases were measured during exploration in the first 30 min of the later session. Lines show linear fits and shading shows 95% confidence intervals. The depth weighted P_BS_ weights each junction choice by 2^6−d^, where *d* is decision node depth. The depth weighted P_BS_ correlation was supported by a two-sided regression with standard errors clustered by animal (BH corrected *p* = 3.7 × 10^−5^).

**Supplementary Table 1.**
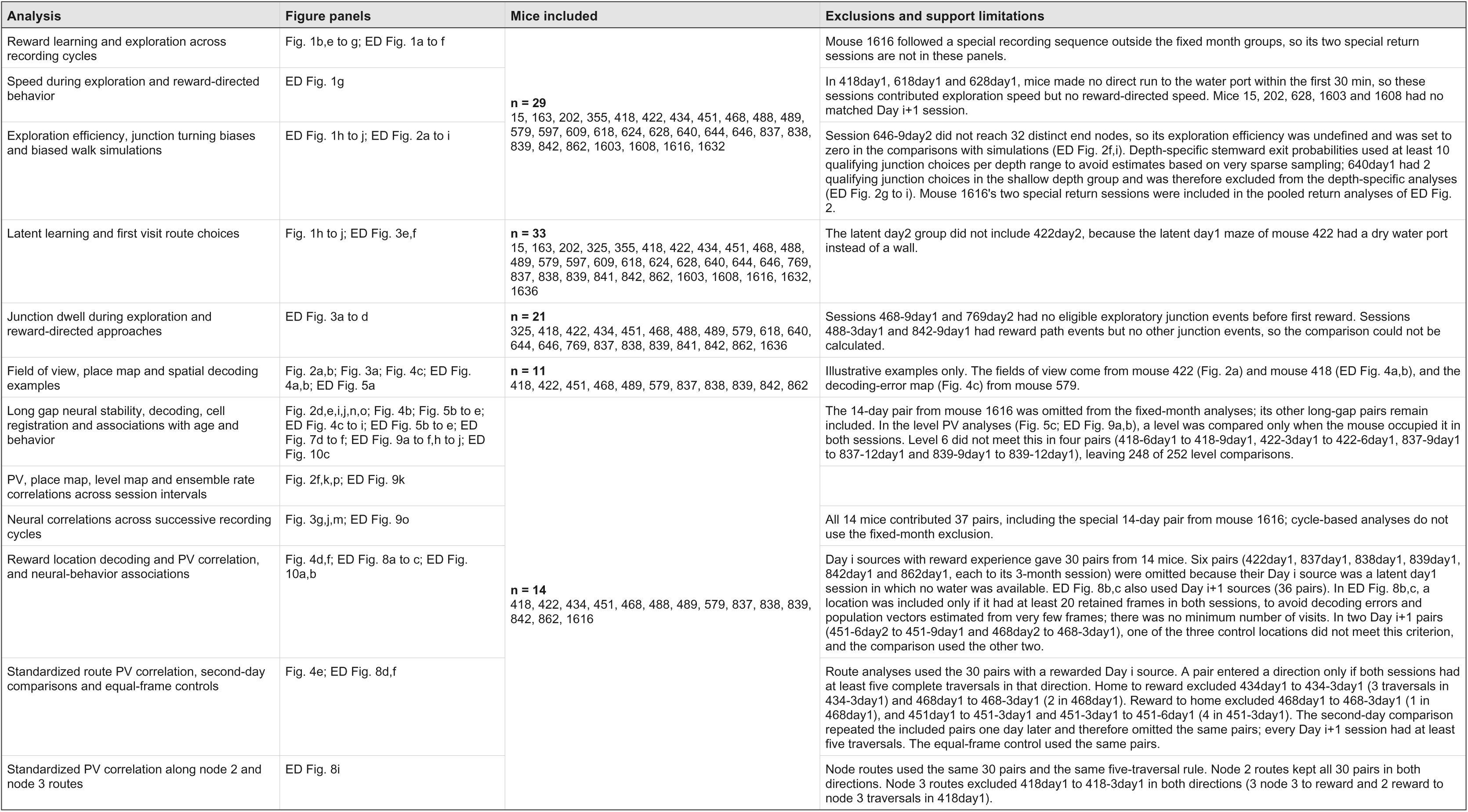
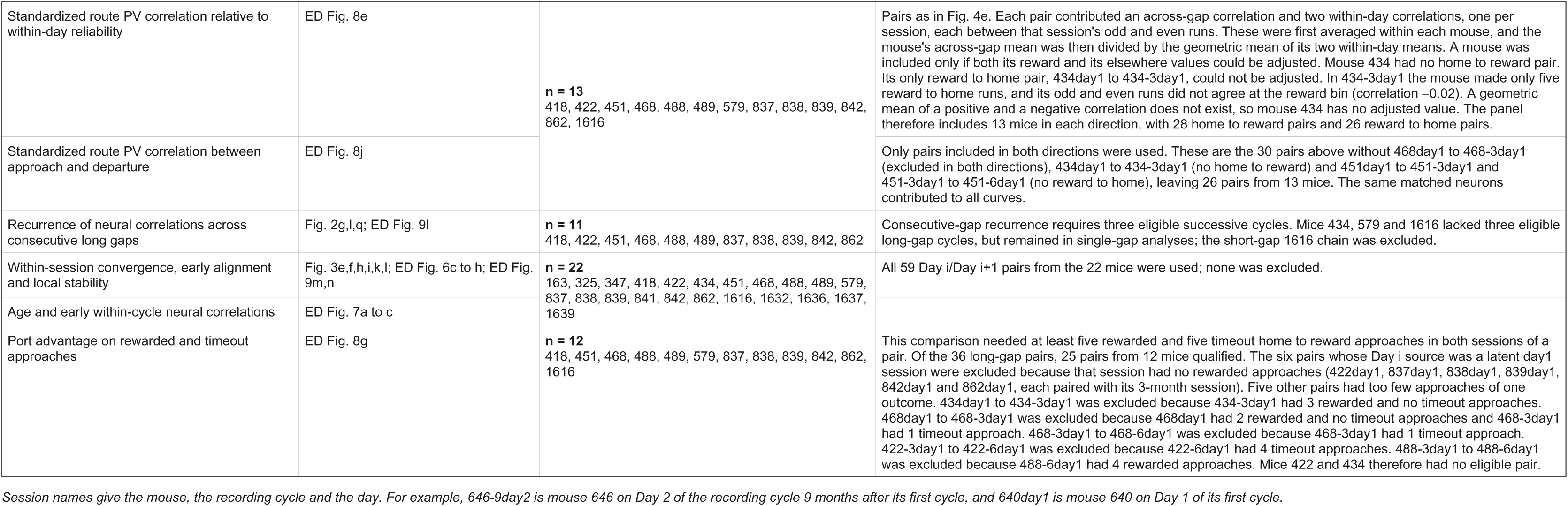
Mouse cohorts by analysis.

**Supplementary Table 2.**
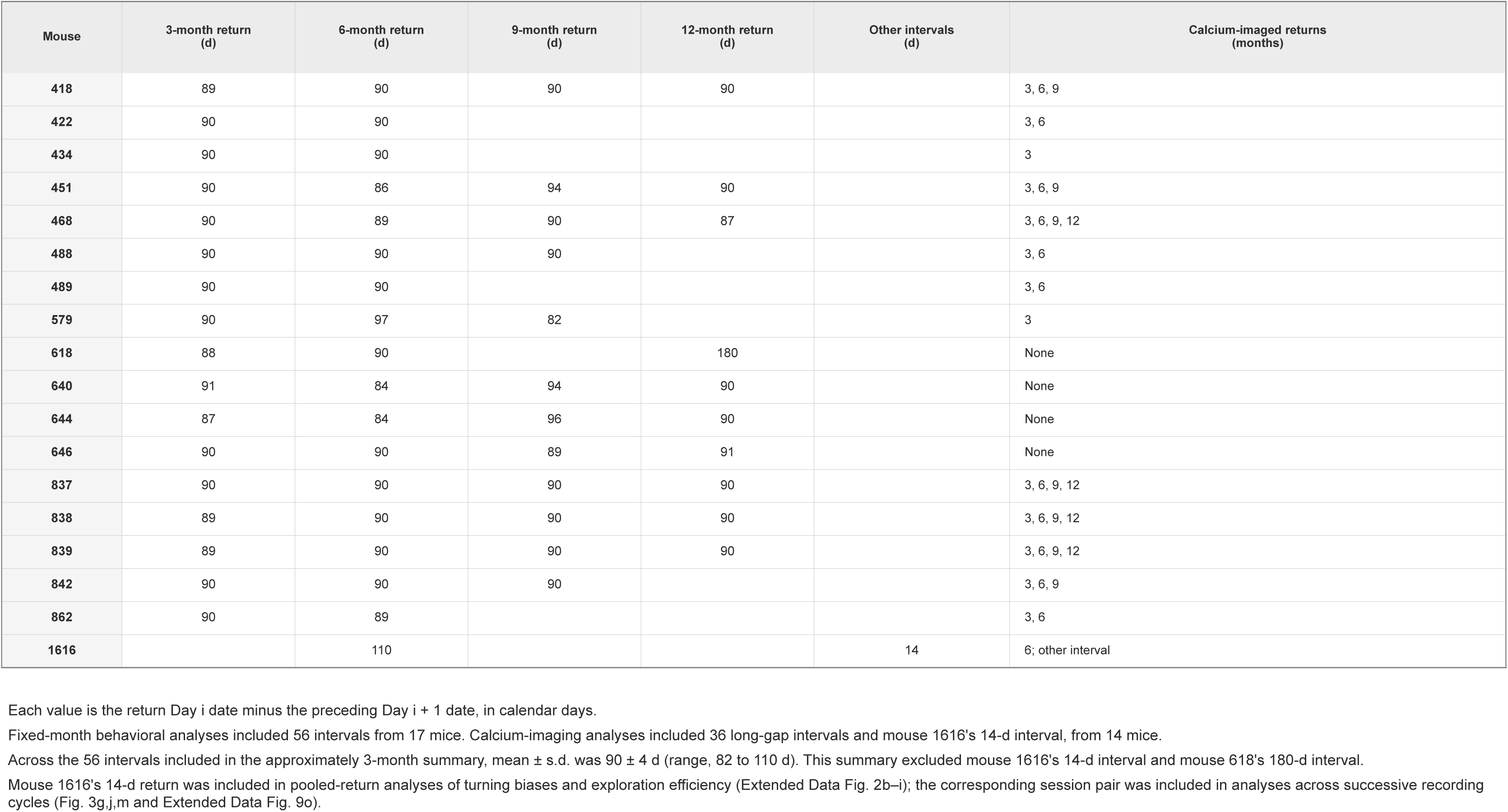
Intervals between successive recording cycles.

| Mouse | 3-month return<br>(d) | 6-month return<br>(d) | 9-month return<br>(d) | 12-month return<br>(d) | Other intervals<br>(d) | Calcium-imaged returns<br>(months) |
|---|---|---|---|---|---|---|
| **418** | 89 | 90 | 90 | 90 |  | 3, 6, 9 |
| **422** | 90 | 90 |  |  |  | 3, 6 |
| **434** | 90 | 90 |  |  |  | 3 |
| **451** | 90 | 86 | 94 | 90 |  | 3, 6, 9 |
| **468** | 90 | 89 | 90 | 87 |  | 3, 6, 9, 12 |
| **488** | 90 | 90 | 90 |  |  | 3, 6 |
| **489** | 90 | 90 |  |  |  | 3, 6 |
| **579** | 90 | 97 | 82 |  |  | 3 |
| **618** | 88 | 90 |  | 180 |  | None |
| **640** | 91 | 84 | 94 | 90 |  | None |
| **644** | 87 | 84 | 96 | 90 |  | None |
| **646** | 90 | 90 | 89 | 91 |  | None |
| **837** | 90 | 90 | 90 | 90 |  | 3, 6, 9, 12 |
| **838** | 89 | 90 | 90 | 90 |  | 3, 6, 9, 12 |
| **839** | 89 | 90 | 90 | 90 |  | 3, 6, 9, 12 |
| **842** | 90 | 90 | 90 |  |  | 3, 6, 9 |
| **862** | 90 | 89 |  |  |  | 3, 6 |
| **1616** |  | 110 |  |  | 14 | 6; other interval |

Each value is the return Day i date minus the preceding Day i + 1 date, in calendar days.
Fixed-month behavioral analyses included 56 intervals from 17 mice. Calcium-imaging analyses included 36 long-gap intervals and mouse 1616's 14-d interval, from 14 mice.
Across the 56 intervals included in the approximately 3-month summary, mean ± s.d. was 90 ± 4 d (range, 82 to 110 d). This summary excluded mouse 1616's 14-d interval and mouse 618's 180-d interval.
Mouse 1616's 14-d return was included in pooled-return analyses of turning biases and exploration efficiency (Extended Data Fig. 2b–i); the corresponding session pair was included in analyses across successive recording cycles (Fig. 3g,j,m and Extended Data Fig. 9o).

**Supplementary Table 3.** Glossary of recurring terms.

| Term | Meaning |
|---|---|
| **Recording cycle** | Two sessions starting 24 h apart, labeled Day i and Day i+1. Successive cycles were separated by a long gap of about 90 days without maze experience. |
| **Day i and Day i+1** | Day i is the first session of a recording cycle: the naive day at initial maze exposure and a return day in later cycles. Day i+1 is the second day of the recording cycle; its session starts 24 h after the Day i session starts. |
| **Latent controls** | Mice that explored without a water port on latent day1. The port was installed for latent day2, when they had prior maze exposure but no prior reward experience. |
| **Session pair** | Two recordings from the same animal compared in an analysis. |
| **Time in maze** | Time spent inside the maze during a session, summed across maze entries. |
| **Maze tile and tile step** | A maze tile is one traversable square of the 15-by-15 maze grid and is also the spatial bin for neural maps and decoding. A tile step is movement between adjacent tiles. |
| **Node, end node and node step** | Nodes are junctions and maze ends (end nodes). A node step joins adjacent nodes and can span several tiles. At a junction, the stem leads toward the entrance and the two branches lead away. |
| **Tree level (depth)** | The number of branch choices from the entrance, ranging from 0 at the entrance to 6 at the end nodes. Locations at the same depth belong to the same level. |
| **Exploration** | Trajectory segments not classified as reward-directed or home-directed travel. Reward-directed travel follows direct routes to the water port with all correct turns in rewarded sessions; home-directed travel follows direct routes home with all correct turns. Exploration efficiency and turning biases are measured during exploration. |
| **Long bout** | A run ending at the water port after more than six consecutive node steps, each reducing the remaining path to it. |
| **First long bout time** | Time in maze before the start of the first long bout. |
| **Exploration efficiency** | 32/*N*<sub>32</sub>, where *N*<sub>32</sub> is the number of end node visits needed to reach 32 distinct end nodes during exploration. Higher values mean fewer repeated visits. |
| **Turning biases** | Probabilities of continuing, reversing or choosing between exits at a junction during exploration. |
| **Deconvolved activity** | The nonnegative estimate of neuronal activity obtained by deconvolving the recorded fluorescence using an exponential-decay model of calcium indicator dynamics. Activity in the neural analyses refers to this estimate. |
| **Frame** | One image in the 25 Hz behavioral recording, corresponding to a 40 ms time step in the aligned data. |
| **Matched neuron (registered neuron)** | A neuron identified as the same cell in different recordings. |
| **Place map and place field** | A place map gives one neuron's mean activity at each maze location. Regions of high activity are its place fields. Place map correlation compares the maps between sessions. |
| **Population vector (PV)** | A vector containing one mean activity value per matched neuron at a given maze location or tree level. PV correlation compares these vectors between sessions. |
| **Ensemble rate correlation** | Correlation across matched neurons between their mean activity in two sessions. It measures whether the relatively active neurons remain relatively active. |
| **Level map** | One neuron's mean activity at each tree level, pooling locations at the same level. A unique peak identifies the preferred level. Level map correlation compares these maps between sessions. |
| **Standardized PV correlation** | PV correlation along a route to or from the water port, after subtracting each neuron's mean activity along the route and dividing by its standard deviation. This compares the neurons' relative preference for each route location. |
| **Neuron-ID shuffle** | Random reassignment of neuron identities in one session, breaking the correspondence between matched cells while preserving their activity. |
| **Decoding error and reward decoding error** | A decoder trained on the earlier (source) session estimates location in the later (target) session. Decoding error is the mean Euclidean distance, in tiles, between decoded and observed locations. Reward decoding error is measured in the tile next to the rewarded end node. |

